# Infection-generated electric field in gut epithelium drives bidirectional migration of macrophages

**DOI:** 10.1101/427294

**Authors:** Yaohui Sun, Brian Reid, Fernando Ferreira, Guillaume Luxardi, Li Ma, Kristen L. Lokken, Kan Zhu, Gege Xu, Yuxin Sun, Volodymyr Ryzhuk, Betty P. Guo, Carlito B. Lebrilla, Emanual M. Maverakis, Alex Mogilner, Min Zhao

**Affiliations:** Department of Dermatology, School of Medicine, University of California, Davis, Sacramento, CA 95817, USA; Courant Institute and Department of Biology, New York University, New York, NY 10012, USA; Departamento de Biologia, Centro de Biologia Molecular e Ambiental (CBMA), Universidade do Minho, Braga 4710, Portugal; Skin and Cosmetic Research Department, Shanghai Skin Disease Hospital, Shanghai, China; Department of Microbiology and Immunology, School of Medicine, University of California, Davis, CA 95616, USA; Department of Chemistry, University of California, Davis, CA 95616, USA; Office of Research, School of Medicine, University of California, Davis, Sacramento, CA 95817, USA

## Abstract

Many bacterial pathogens hijack macrophages to egress from the port of entry to the lymphatic/blood-stream, causing dissemination of life-threatening infections. However, the underlying mechanisms are not well understood. Here, we report that *Salmonella* infection generates directional electric fields (EF) in the follicle-associated epithelium of mouse cecum. *In vitro* application of an EF, mimicking the infection-generated electric field (IGEF), induces directional migration of primary mouse macrophages to the anode, which is reversed to the cathode upon *Salmonella* infection. This infection-dependent directional switch is independent of the *Salmonella* pathogenicity island 1 (SPI-1) type III secretion system. The switch is accompanied by a reduction of sialic acids on glycosylated surface components during phagocytosis of bacteria, which is absent in macrophages challenged by microspheres. Moreover, enzymatic cleavage of terminally exposed sialic acids reduces macrophage surface negativity and severely impairs directional migration of macrophages in response to EF. Based on these findings, we propose that macrophages are attracted to the site of infection by a combination of chemotaxis and galvanotaxis; after phagocytosis of bacteria, surface electrical properties of the macrophage change, and galvanotaxis directs the cells away from the site of infection.

**Abbreviations:** CFU, colony-forming unit; Con A, Concanavalin A; EF, electric field; FAE, follicle-associated epithelium; GNL, Galanthus Nivalis lectin; IGEF, infection-generated electric field; Ji, electric current density; MAL-2, Maackia Amurensis lectin II; MLN, mesenteric lymph node; MOI, multiplicity of infection; nMFI, normalized mean fluorescence intensity; RCA-1, Ricinus Communis Agglutinin I; SNA, Sambucus Nigra lectin; *S*. Typhimurium, *Salmonella enterica* serotype Typhimurium; SPI-1, *Salmonella* pathogenicity island 1; PDMS, polydimethylsiloxane; TEP, trans-epithelial potential difference; TLR, Toll-like receptors; WGEF, wound-generated electric field

## Introduction

Common bacterial pathogens such as *Salmonella, Shigella* and *Yersinia* spp. invade the gut epithelial barrier, preferentially by targeting the relatively small number of M cells located in the follicle-associated epithelium (FAE) (1, 2). Disruption of epithelial integrity releases chemokines that attract immune cells such as neutrophils and macrophages—a process known as chemotaxis (3-6). Subsequent phagocytosis and clearance of these pathogens by immune cells usually stops the infection. However, some of these bacterial pathogens have developed strategies, such as the type III secretion systems in *Salmonella* spp. (7-11), to evade macrophage killing and survive inside the macrophage (12-15), an environment in which the pathogen is hidden from the immune system. Survival within the macrophage allows the pathogen to spread from its entry site to the spleen, liver, bone marrow, and other organs via the blood-stream, resulting in life-threatening consequences (16-18). While chemotaxis can explain how macrophages reach an infected site, it cannot explain how macrophages harboring pathogens escape from the bacterial entry site to reach the lymphatic drainage/blood-stream, a critical initial step in the dissemination process that is understudied and poorly understood.

Bioelectric signals have been implicated in development (19-22), wound healing (23-25) and regeneration (26, 27). For example, a wound collapses the transepithelial potential (TEP) difference of an intact epithelial barrier, generating laterally oriented endogenous electric fields (EF) of up to 1.4 V cm^−1^, as well as local electric current densities (J_I_) of several μA cm^−2^. These bioelectric phenomena actively control wound healing in the skin and cornea (24, 28); however, they are extremely challenging to study in the gut epithelium due to limited accessibility, and have never been characterized *in vivo* during an active infection. Nonetheless, it is generally appreciated that EF on this scale can drive directional cell migration—a process known as electrotaxis or galvanotaxis (20). Many cell types, regardless of their origin, respond to exogenous EF by directional migration toward the cathode (24, 29), while others, *e.g*., human keratinocytes (HaCat cells) and bone marrow mesenchymal stem cells, migrate toward the anode (30). Macrophages and lymphocytes also undergo galvanotaxis *in vitro* and *in vivo* (31-33).

In the present study, we have developed an *ex vivo* mouse cecum model of *Salmonella* infection that enables bioelectricity measurement. We report that *Salmonella* infection generates directional EF at the bacterial entry sites that can recruit macrophages by galvanotaxis. We demonstrate that primary macrophages reverse galvanotaxis direction upon *Salmonella* infection by modifying their surface glycan composition, which reduces the negative electric charge on the surface. This directional switch is independent of the *Salmonella* pathogenicity island 1 (SPI-1) type III secretion system, a major virulence determinant responsible for *Salmonella* invasion. Instead, it may require certain glycosidases that are widely conserved in *Salmonella* spp., as cleavage of surface-exposed sialic acids with a potent neuraminidase caused severe defects of macrophage galvanotaxis.

## Results

### Development of an *ex vivo* mouse cecum model of *Salmonella* infection for bioelectric characterization

The mouse is an ideal organism for understanding human infectious diseases and is widely used to study bacterial pathogenesis and mucosal immunity of enteropathogenic bacteria (34-39). Previously, we have successfully measured bioelectric currents in various tissue and organ cultures using the non-invasive vibrating probe (40-43); however, measuring bioelectric currents in the mouse small intestinal epithelium is challenging due to limited accessibility (44). Therefore, we developed a new *ex vivo* cecum model for measuring bioelectric activity in the gut epithelium (**S1 Fig.**). This model is based on the well-established mouse typhoid model, where mice orally infected with *S*. Typhimurium develop disseminated infection that mimics human typhoid disease (35). Although the ileum is the most commonly targeted organ to study pathogen-host interactions *in vivo*, it is too small for the bioelectrical measurements in our *ex vivo* experimental setting. Instead, the size of the mouse cecum is anatomically optimal (45). *Salmonella* invades the cecum in mice, and causes acute appendicitis in humans (46). Furthermore, we can easily identify the FAE under a dissecting microscope, as we found that 90% of C57BL/6 mice have only one Peyer’s patch around the blind-end apex, containing 6-11 lymphatic follicles (**S3B Fig.**).

### Active bioelectricity pervades the FAE in the healthy murine cecum

Using microelectrodes (27) in the *ex vivo* mouse cecum model, we measured a TEP of up to 15 mV, lumen-positive (**Fig. 1A**) in uninfected mice. Notably, the TEP in the FAE was significantly smaller than that of the surrounding villi (**Fig. 1B**). Next, using a vibrating probe to measure the J_I_ close to the gut surface, we detected outward currents at the FAE and inward currents at the villus epithelium, with a magnitude of around 1 μA cm^−2^ (**Fig. 1C**). Such currents were not detected in the serosal epithelium, despite the presence of a TEP, nor in formalin-fixed epithelia (**Fig. 1D**), suggesting the existence of active bioelectricity restricted to the mucosal epithelium.

**Fig. 1.**
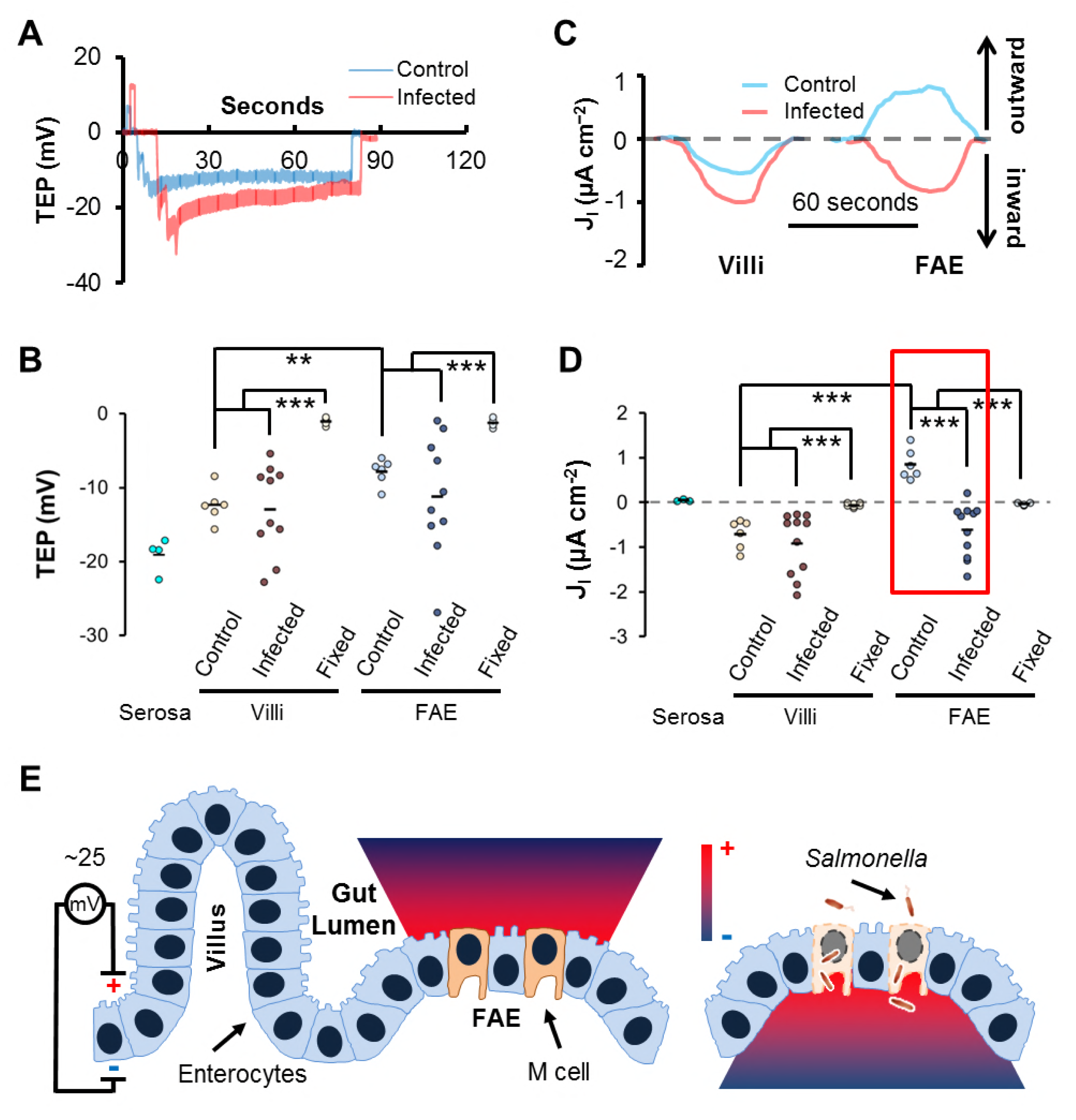
Infection-generated electric field (IGEF) at gut epithelium. (A) Representative TEP show stepwise increase (more negative inside) at the FAE after oral *Salmonella* infection (red), compared to an uninfected control (blue). (B) Summary of TEP in control mice or mice at 16 to 40h post of infection (POI). Each dot represents an average of 3-5 follicles or other epithelia of each mouse. Serosal epithelia and formalin-fixed mouse ceca were served as controls. ** *P* < 0.01 by unpaired Student’s t-test. *** *P* < 0.001 by unpaired Student’s *t*-test. (C) Representative J_I_ measured by vibrating probe. *Salmonella* infection (red) induces a reversed inward current in FAE compared to an outward current in the uninfected control mice (blue). (D) Summary of peak J_I_ Each dot represents an average of 3-5 follicles or other epithelia of each mouse. Serosal epithelia and formalin-fixed mouse ceca were served as controls. *** *P* < 0.001 by unpaired Student’s *t*-test. The red box highlights *Salmonella* infection-generated current reversal at the FAE. (E) The diagram depicts IGEF generated by *Salmonella* infection at the gut epithelium in a mouse model of human typhoid fever. TEP up to 25 mV, lumen-positive, occur across a single layer of tightly sealed gut epithelium and drive micro-ionic currents running from the epithelial surface to the lumen (color-coded gradient). *Salmonella* invades and breaks epithelial integrity, preferentially at the FAE, which generates ionic currents running from the breached epithelium into the deep intestinal wall (color-coded gradient).

### *Salmonella* infection generates bioelectric fields at the FAE of its entry site

In mice intragastrically challenged with *Salmonella enterica* serotype Typhimurium (*S*. Typhimurium), the peak J_I_ remarkably reversed to become inward in FAE compared to control mice, while that of the villus epithelium increased non-significantly (**Fig. 1C and D**). Consistent with this finding, we also detected high variation in TEP from individual *S*. Typhimurium-infected mice (up to 25 mV), still lumen-positive (**Fig. 1A and B**), which has also been observed in chemically induced colitis in rats (47). More importantly, penetrating electrodes through disrupted FAE, up to 200 μm in depth, revealed a stepwise increase in TEP (**Fig. 1A and S2 Fig.**), suggesting the existence of an electric potential gradient generated by the *S*. Typhimurium infection.

Bacterial invasion and subsequent dissemination to mesenteric lymph nodes (MLN) and spleen were verified by determination of colony-forming units (CFU) (**S4A Fig.**), and disruption of FAE was assessed by histological staining (**S4B Fig.**). *S*. Typhimurium was undetectable in some of the MLN and spleens, indicating that some of the FAE and surrounding villi were either uninfected or mildly infected, which explains the wide distribution of TEP and J_I_ measured in the *S*. Typhimurium-challenged mice. Based on these results, we coined the term *infection-generated* electric field (IGEF) (**Fig. 1E**) to distinguish it from a *wound-generated* electric field (WGEF) (25).

### *Salmonella* reverses the directional migration of IGEF-guided macrophage galvanotaxis *in vitro*

We next sought to understand the role of the IGEF during *S*. Typhimurium infection. In response to an exogenous EF, tuned to mimic the *in vivo* IGEF (mathematics in Materials and Methods), the mouse peritoneal-derived macrophages demonstrated robust unidirectional migration to the anode (**S1 Movie**)—replicating the active biological process known as galvanotaxis (20, 25). This unidirectional migration was verified by immunostaining showing that nearly 100% macrophages were polarized to the anode with a distinct morphology characterized by a leading pseudopodium of dense actin meshwork and a rearward uropod (**Fig. 2F**). However, upon challenge with *S*. Typhimurium IR715, the average directionality of EF-induced galvanotaxis decreased significantly, with approximately 41% of the macrophages reversing their migratory direction from the anode to the cathode (**Fig. 2, and S2 Movie**). This migration reversal phenotype was robustly reproduced in bone marrow-derived macrophages challenged by three virulent *S*. Typhimurium strains: IR715, SL1344 and LT2 (**S5 Fig.**). If the cells had simply stopped sensing the EF, then we would have observed random migration, as was the case for control macrophages not subjected to an EF (**Fig. 2E**). The observed change in migration pattern (**Fig. 2E**) can be attributed to *S*. Typhimurium infection for a variety of reasons. First, a gentamycin protection assay confirmed the presence of intracellular bacteria (**Fig. 2D**); second, flow cytometry demonstrated a high *S*. Typhimurium infection rate (~53%) (**S7 Fig.**); and third, high-resolution confocal microscopy revealed that macrophages containing intracellular *S*. Typhimurium switched polarity to the cathode (**Fig. 2G**). Based on these findings, we conclude that *Salmonella-containing* macrophages respond to galvanotaxis stimuli by reversing their primary directional migration.

**Fig. 2.**
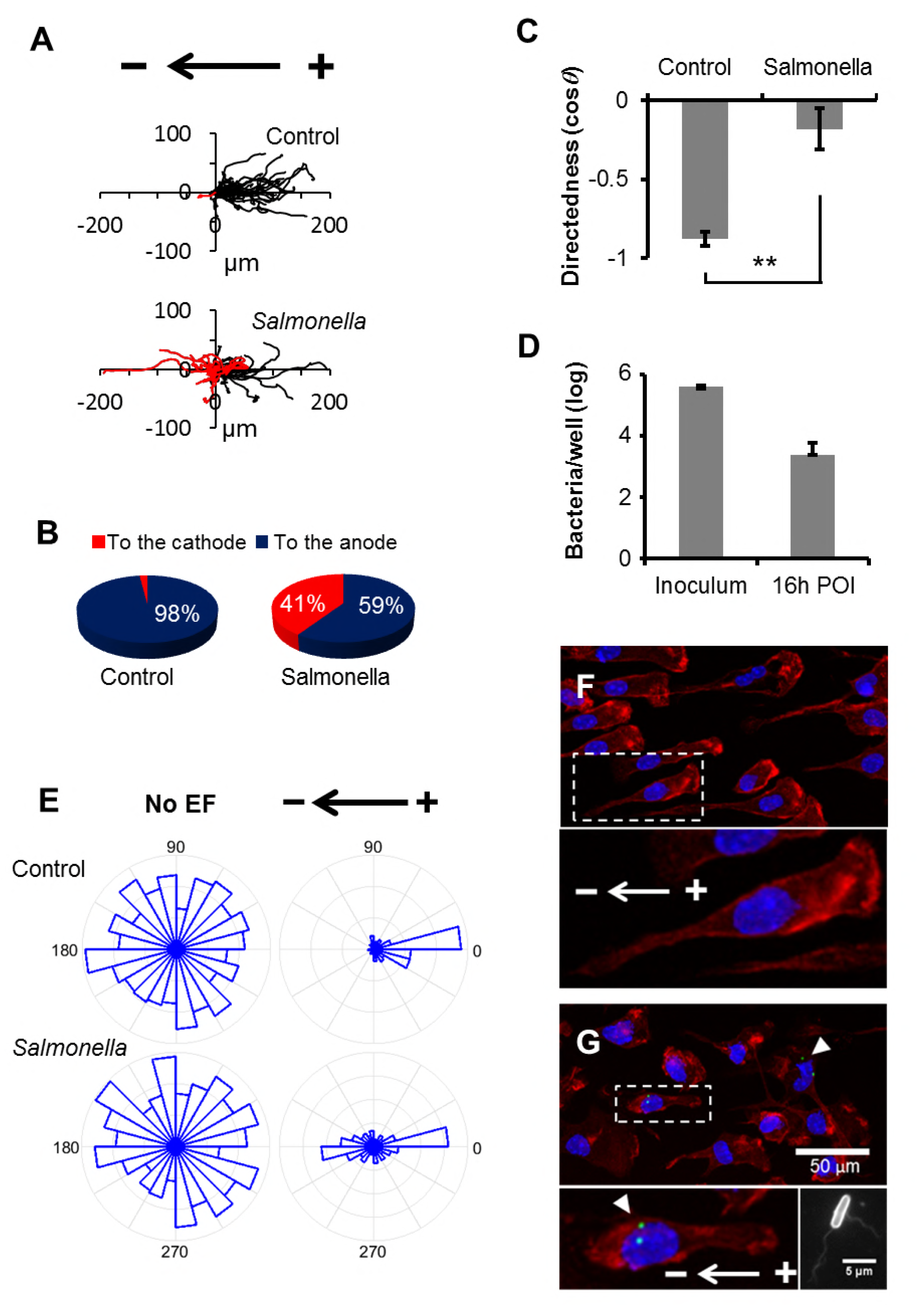
*Salmonella* infection switches macrophage galvanotaxis to the cathode. (A) Trajectories of peritoneal macrophages exposed to an IGEF-mimetic EF of 4 V cm^−1^ in the indicated orientation before or at 16 h POI. Each line represents one cell’s trajectory. Duration was 3 h with time zero placed in the center. (B) Pie charts show percentage of cells migrating to the cathode (red) or to the anode (blue) as demonstrated in panel A. (C) Quantification of overall directionality as directedness. Directedness is calculated as cosine theta (*θ*), where *θ* is the angle of each cell traveled corresponding to the applied EF field (positive, to the cathode; negative, to the anode). Data from a representative of multiple independent experiments. ** *P* < 0.01 by unpaired Student’s *t*-test. (D) Intracellular bacteria were quantified per gentamycin protection assay in 24-well plates and normalized by inoculum. (E) Rose plots show random migration of macrophages with no EF, unidirectional migration (to the anode) of control macrophages in response to an EF of 4 V cm^−1^, and bidirectional migration (either to the anode or the cathode) of cells infected with *Salmonella* exposed to the same EF. (F) A representative confocal image shows that macrophages were polarized to the anode with a characteristic morphology: massive actin meshwork labeled by Alexa Fluor 555 Phalloidin (red) in the front and a uropod at the rear. Nuclei were labeled by Hoechst 33342 (blue). Bottom panel shows enlargement of the checked area in the indicated field. (G) A representative confocal image showing that macrophages with intracellular *Salmonella* (white arrowheads) were polarized to the cathode. Intracellular *Salmonella* were detected by a specific antibody that recognizes both cell body and flagellae (bottom right panel) and stained with a secondary antibody conjugated with Alexa Fluor 488 (green). Actin and nuclei were stained as in panel F. Bar, 50 μm. Bottom left panel shows enlargement of the checked area in the indicated field.

### Infection-dependent directional switch of macrophage galvanotaxis is SPI-1independent

Macrophages are professional phagocytes that uptake a broad range of substances; meanwhile, pathogenic *Salmonella* has developed several virulence means to evade macrophage killing, with SPI-1 being the major virulence factor responsible for colonization and invasion (9, 48). To better understand whether the directional reversal is phagocytosis-dependent or SPI-1-specific, we monitored galvanotaxis under identical conditions in macrophages challenged with: i) fluorescently labeled microspheres similar in size to bacteria, ii) *S*. Typhimurium constitutively expressing mCherry, and iii) a GFP-expressing *ΔinvA* mutant that lacks a functional SPI-1 (unable to inject its effectors into cells) (49) (**Fig. 3A**). Macrophages challenged with microspheres exhibited migratory behavior similar to that of controls, *i.e*., unidirectional migration to the anode (**Fig. 3B and S3 Movie**). Surprisingly, macrophages challenged with Δ*invA* migrated with a significantly decreased overall directionality close to that of macrophages infected with wildtype (WT) *Salmonella* (**Fig. 3B**), even though the number of intracellular mutants was indeed lower than that of the WT (**Fig. 3C**). Time-lapse recording captured a marked opposing directional migration pattern: macrophages containing microspheres migrated to the anode and macrophages with either WT or Δ*invA* bacteria inside migrated to the cathode (**Fig. 3D and E, and S3 Movie**), further confirming that the observed directional switch was *Salmonella* infection-dependent. Importantly, there were similar phagocytosis/infection rates between the macrophages challenged with microspheres or *Salmonella* in the given MOI (**S6A and B Fig**.), as verified by flow cytometry (**S7 Fig.**). Cells containing no or variable microspheres migrated to the anode exclusively (**S6C and D Fig**.), ruling out mechanisms solely based on phagocytosis. The fact that cells containing either WT or Δ*invA* migrated to the cathode suggests that the SPI-1 type III secretion system is not required, or at least is insufficient for the directional switch. This data is in accordance with previous studies that identified an SPI-1-independent pathway contributing to early dissemination of *S*. Typhimurium in the mouse typhoid model (16). We therefore argue for the existence of a general infection-dependent mechanism that involves phagocytosis and subsequent interplay between the macrophages and internalized bacterial pathogens.

**Fig. 3.**
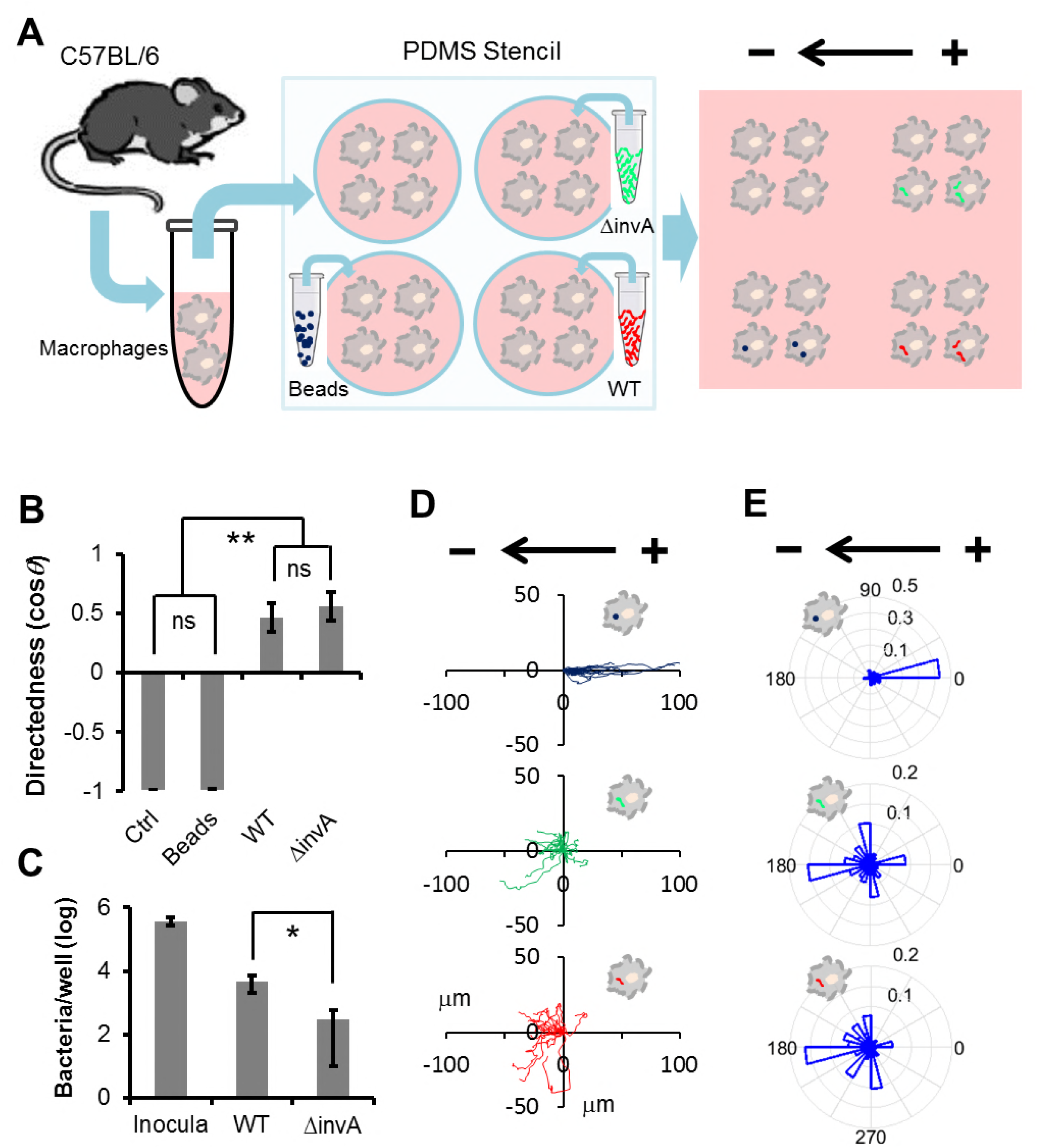
Directional switch mediated by *Salmonella* infection in macrophage galvanotaxis is phagocytosis-independent, and SPI-1-independent. (A) Schematic of the experimental design. Freshly differentiated mouse bone marrow-derived macrophages were seeded in wells engineered with a PDMS stencil. Adhered macrophages were challenged with either blue fluorescent beads or wildtype *Salmonella* (WT) expressing mCherry or SPI-1 mutants (Δ*inνΛ*) expressing GFP. Subsequent migration of macrophages was monitored in the same galvanotaxis chamber under identical conditions, after removal of the PDMS stencil. (B) Directedness of macrophages under different challenge conditions. ns, non-significant difference between the control macrophages and the macrophages challenged with beads. ** *P* < 0.01, compared to the control or beads group by unpaired Student’s *t*-test. (C) Quantification of intracellular bacteria by gentamycin protection assay. Representative data presented as log CFU per well, normalized to each inoculum. Bar in S.E. from triplicate wells. * *P* < 0.05 by unpaired Student’s *t*-test. (D) Trajectories of macrophages bearing beads (blue) or WT (red) or *ΔinvA* (green) exposed to an EF of 4 V cm ^−1^ in the indicated orientation. Duration was 3 h. See also S3 Movie. (E) Rose plots show opposite galvanotactic behaviors of the macrophages bearing beads (to the anode) or the macrophages containing WT or Δ*invA* (to the cathode) in response to an EF of 4 V cm^−1^ in the indicated orientation.

### *Salmonella* but not microspheres decreases surface-exposed sialic acids in macrophages

Charged cell-surface components are critical for EF-induced motility in 3H3 cells (50) and have been implicated in electro-osmosis of concanavalin A (Con A) binding receptors on the surface of myotomal spheres (51). We hypothesized that the directional switch of macrophage galvanotaxis could result from bacteria-induced modifications to the charged components of the cell surface, which would not occur following microsphere challenge (**Fig. 4A**). To this end, we screened *Salmonella-infected* and control macrophages against a panel of fluorescently labeled lectins (glycan-binding proteins), capable of detecting charged and uncharged cell surface glycans (**S3 Table**). The normalized mean fluorescent intensity of Maackia Amurensis Lectin II (MAL-2), a lectin that recognizes pathogen-binding sialic acids, was significantly decreased in macrophages infected by *Salmonella* but not in those carrying microspheres (**Fig. 4B to D**). Marked Galanthus Nivalis Lectin (GNL)- and Con A-binding aggregates were visible within macrophages infected by *Salmonella* (**S8 and S9 Fig.**), raising the possibility that the decrease in MAL-2-binding sialic acids could be the result of bacterial internalization and subsequent degradation.

**Fig. 4.**
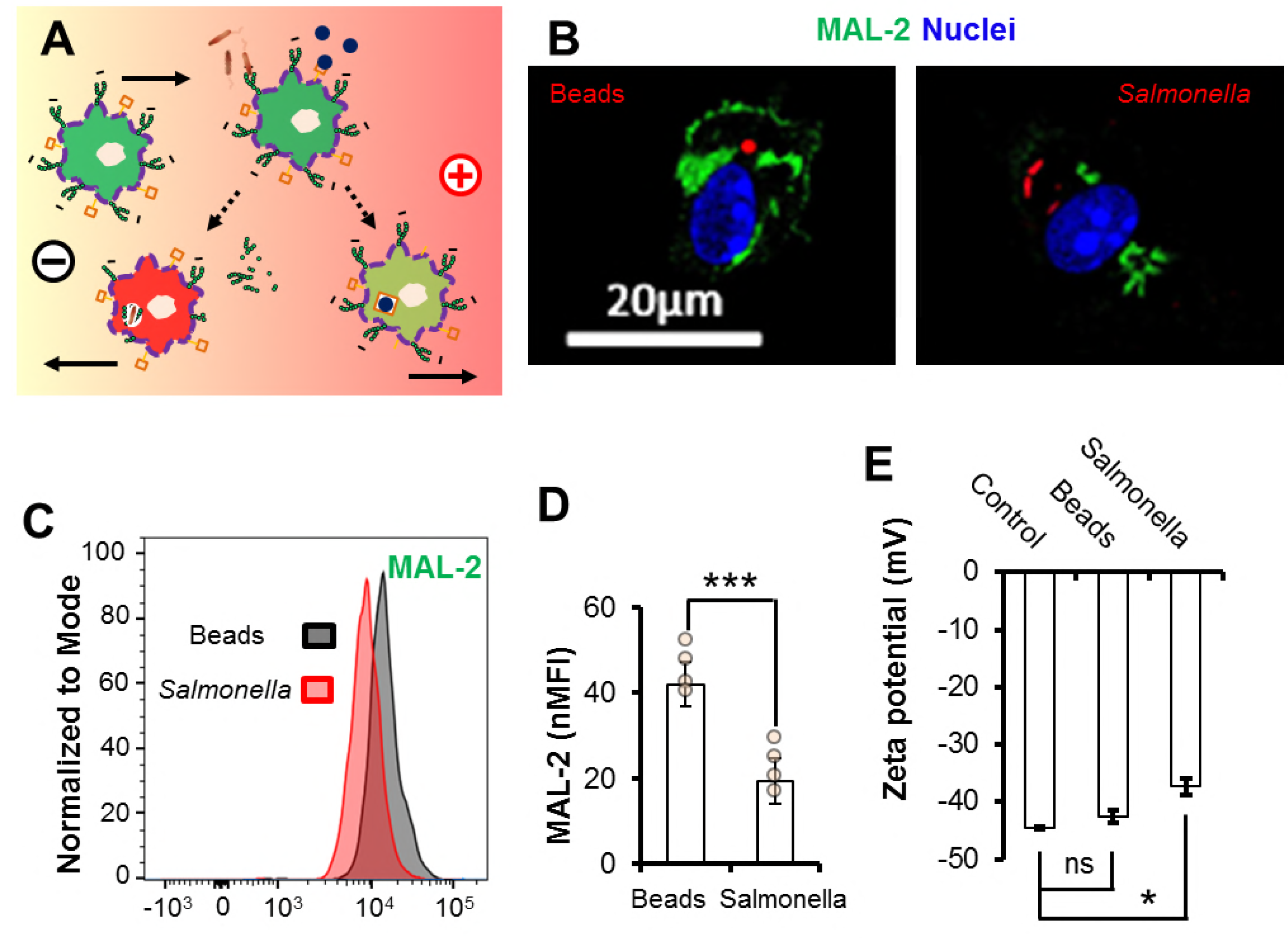
Decrease of sialylated surface components and reduction of negative charges mediate *Salmonella* infection-dependent directional switch in macrophage galvanotaxis. (A) Schematic showing the simplified hypothesis of galvanotaxis in macrophages and a directional switch modulated by *Salmonella* infection. Primary macrophages with sialylated glycoproteins or lipids (small green circles), migrate to the anode. Activated macrophages with reduced negativity, either through enzymatic activities of the *Salmonella* or internalization/degradation, switch direction to the cathode while macrophages that phagocytosed beads via non-sialylated surface components (brown squares), still migrate to the anode. (B) Representative confocal images of surface MAL-2 (green) of macrophages challenged with 1 μm diameter red fluorescent microspheres or *S*. Typhimurium constitutively expressing mCherry (red) at 16h POI. Nuclei were counterstained with DAPI (blue). Bar, 20 μm. (C) Representative flow cytograms and (D) independent data of standardized MAL-2 fluorescence (green) intensity of macrophages treated as in panel B. ns, non-significant. *** *P* < 0.001 by Student’s *t*-test. (E) Zeta potential of control macrophages or macrophages challenged with beads or *S*. Typhimurium at 16 h POI. Data were quantified from a representative of three independent experiments. ns, non-significant. * *P* < 0.05 by Student’s *t*-test.

### Cleavage of surface-exposed sialic acids impairs macrophage galvanotaxis via zeta potential

Sialylated cell surface molecules are negatively charged, creating an electronegative zeta potential (52). Using an electrophoretic light scattering technique, we determined the zeta potential of macrophages under various conditions. The negative zeta potential of macrophages infected by *S*. Typhimurium was significantly reduced, *i.e*., less negative than that of control macrophages (*P* < 0.05) (**Fig. 4E**). By contrast, macrophages challenged with microspheres showed a non-significant zeta potential change compared to that of control macrophages (**Fig. 4E**), suggesting that the decrease in surface-exposed sialic acids and subsequent reduction of surface negativity is mediated by active bacterial product(s) (53).

If a decrease of the negatively charged sialic acids of the surface glycoproteins is critical for the directional switch in macrophage galvanotaxis, then cells with their sialic acids enzymatically removed should exhibit a switch or at least a defect in directional migration when exposed to an EF. To test this, we treated freshly isolated mouse macrophages with a potent neuraminidase (an enzyme that cleaves terminal sialic acid residues from surface-exposed glycoproteins) (54). Cleavage of sialic acids following enzymatic treatment was confirmed by flow cytometry and confocal microscopy (**Fig. 5A to C**). As predicted, the zeta potentials of neuraminidase-treated macrophages were significantly reduced (**Fig. 5D**). These cells also lost anodal migration compared to control macrophages monitored in parallel (**Fig. 5E and F, and S4 Movie**). In fact, actin/lectin staining and morphological quantification showed that nearly 12% of the cells were polarized to the cathode (**Fig. 5G**). To further determine the importance of surface negativity, we incubated macrophages in medium at pH 5.8 that markedly reduced the zeta potential (**S11A Fig.**). Similar to the neuraminidase treatment, galvanotaxis of macrophages under acidic conditions was significantly impaired, resulting in 14% of the macrophages reversing their polarity to the cathode (**S11B to D Fig. and S5 Movie**).

**Fig. 5.**
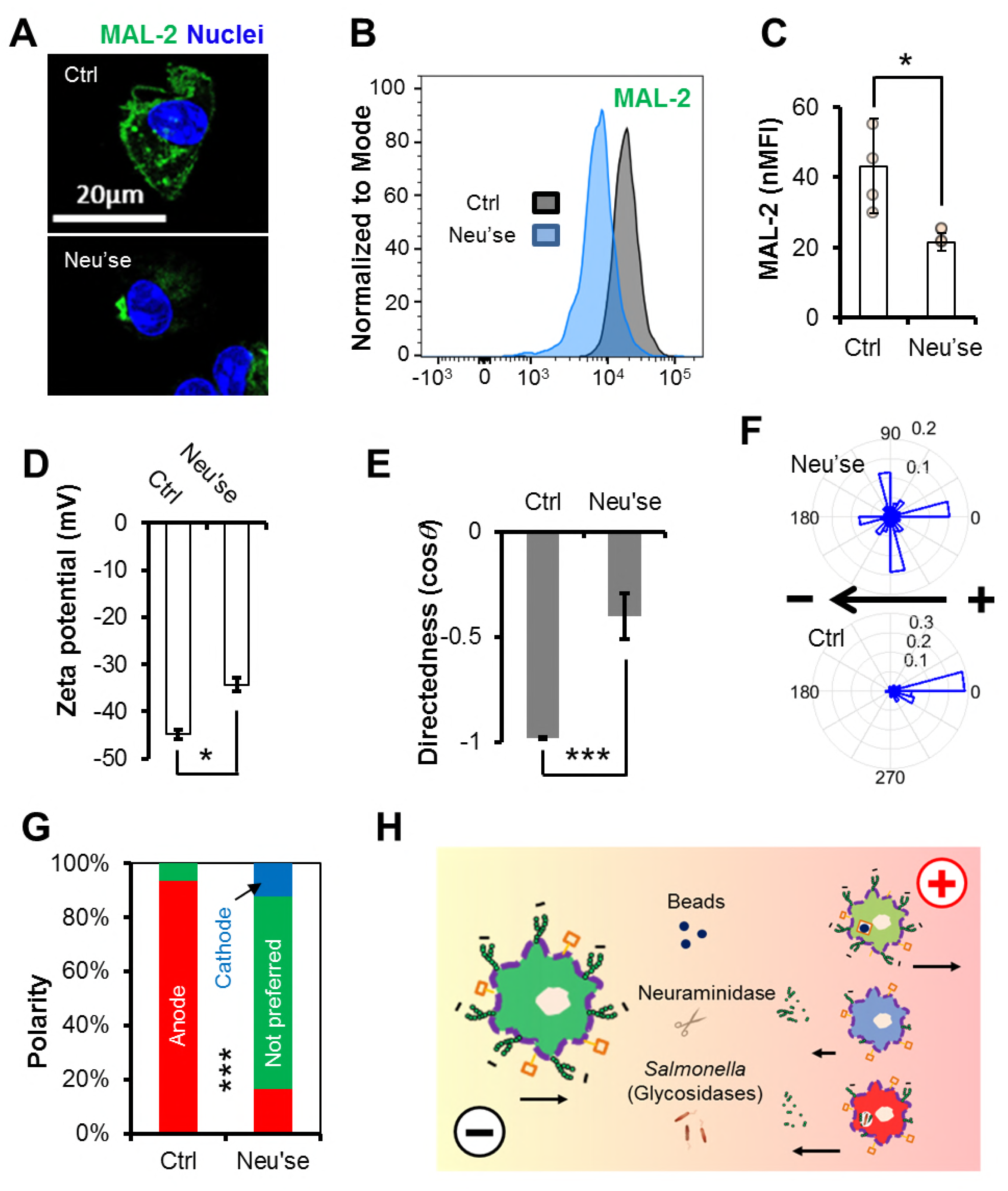
Cleavage of negatively-charged sialic acids impairs macrophage galvanotaxis. (A) Representative confocal images of surface MAL-2 (green) of macrophages incubated with 0 or 100 mU ml^−1^ neuraminidase for 30 minutes. Nuclei were counterstained with DAPI (blue). Bar, 20 μm. (B) Representative flow cytograms and (C) independent data of standardized MAL-2 fluorescence (green) intensity of macrophages incubated with 0 or 100 mU ml^−1^ neuraminidase for 30 minutes. * *P* < 0.05 by Student’s *t*-test. (D) Zeta potential of macrophages incubated with 0 or 100 mU ml^−1^ neuraminidase for 30 minutes. Data were quantified from a representative of three independent experiments. ns, non-significant. * *P* < 0.05 by Student’s *t*-test. (E) Directedness, (F) rose plots and (G) polarity of macrophages treated with or without neuraminidase, followed by 3 h exposure to an EF of 4 V cm^−1^. Data were quantified from a representative of four independent experiments. *** *P* < 0.001 by Student’s *t*-test (Panel E). *** *P* < 0.001 by χ^2^ test (Panel G). (H) Proposed model of *Salmonella* infection-dependent directional switch in macrophage galvanotaxis. Upon exposure to an IGEF-like EF (marked gradient), negatively charged (sign with small green circles) macrophages undergo robust directional migration to the anode. Phagocytosing *Salmonella* reduces surface negativity of the macrophage through catalytic activities of certain glycosidases, as exemplified by the cleavage of sialic acids with a neuraminidase (scissors). Consequently, activated macrophages can either undergo defective directional migration or switch direction to the cathode. Macrophages that phagocytosed beads through binding of non-sialylated surface components (brown squares) still migrate to the anode. Arrows indicate both direction and strength of macrophage galvanotaxis.

## Discussion

It has been well-appreciated that mammalian intestinal mucosa maintains a large transmucosal potential difference (55, 56). In humans, that potential is up to 12 mV, lumen-negative in the fasting jejunum and ileum (57, 58). Rats, mice, chickens and fish all maintain a TEP of up to 5 mV in their intestinal epithelium, as measured with Ussing chambers (59-63); however, these measurements provide no spatiotemporal information (44). The TEP of control murine ceca (up to 15 mV) that we have measured directly with glass electrodes under microscopic resolution were well within the physiological range described above; however, it differed spatially, *i.e*., the TEP of FAE was significantly lower than the TEP of the surrounding villus epithelium (**Fig. 1B**). Using an ultrasensitive vibrating probe, we further detected ionic currents that run in opposite directions between the FAE and surrounding villi (**Fig. 1D**). Together, these findings unveil lateral voltage gradients and/or constant current loops running between these two structurally and functionally distinct epithelia. In the normal gut scenario, such a bioelectrical landscape may prevent commensals from accidentally entering the FAE and enable the pathogens to specifically target M cells via bacterial galvanotaxis (64).

One of the major findings of this work is the discovery of infection-generated electric field (IGEF). *Salmonella* invades the intestinal epithelium, preferentially by targeting M cells located at an FAE (1, 65). If a wound disrupting an epithelium can generate a steady EF, one would envisage that similar EF must be produced at the *Salmonella* entry site because of the breakage of epithelial integrity and subsequent short-circuit of the transmucosal potential difference. This is indeed the case in our model. We demonstrate, for the first time, that *Salmonella* infection generated a steady EF (up to 5 V cm^−1^, provided an epithelial thickness of 50 μm) that drives minute directional electric currents, running from the breached FAE into the deep intestinal wall in a step-wise manner (**Fig. 1E**). Like the injury currents reported by Sawyer and colleagues in early publications (66, 67), IGEF-driving ionic currents could affect small blood vessels in the intestinal wall and mesentery to cause a transvascular potential drop or reversal, resulting in two possible consequences: an intravascular occlusion that may benefit transendothelial penetration of immune cells (e.g., monocytes, leukocytes), and/or the creation of a galvanotactic route between the infected epithelium and the electrically impacted vessels (68).

While IGEF may provide a guidance cue for the enterocytes or even local stem cells, contributing to the repair process of damaged epithelium, the major focus of this work is rather to investigate its bio-pathological role in the systemic *Salmonella* infection, specifically during the initiation of macrophage-driven dissemination. Numerous studies have shown that applying EF *in vitro* can direct macrophage galvanotaxis to the anode (31-33). We confirmed this phenotype and further show that the robust anodal migration of macrophages in response to IGEF-like electric fields was not triggered by phagocytosis itself, since phagocytosed microspheres did not alter macrophage galvanotaxis (**Fig. 3 and S3 Movie**). Beyond that, the most dramatic finding of this work is the *Salmonella* infection-dependent directional switch, which we have demonstrated in both peritoneal and bone marrow-derived macrophages, and reproduced by infection with three pathogenic *Salmonella* strains.

How does *Salmonella* infection instruct macrophages to reverse EF-directed migration? Firstly, it is important to note that infecting macrophages with a SPI-1 mutant resulted in a directional switch similar to that of wildtype, with nearly all the macrophages containing live fluorescent protein-expressing bacteria migrating to the cathode (**Fig. 3 and S3 Movie**). This suggests a general mechanism, independent of this major virulence factor, even though other specific factor(s) may still be involved (69, 70). Previously reported effects have suggested that negatively charged surface glycan moieties are critical for EF-induced cell motility and polarization (51), which are consistent with our data, and provide a long sought-after mechanism of action. Since macrophages challenged with microspheres did not show a significantly reduced zeta potential (**Fig. 4E**), it is likely that the decrease in surface-exposed sialic acids and reduction of surface negativity is mediated by active bacterial product(s), rather than by metabolic changes in the host itself. It is also important to note that although both *Salmonella* infection and neuraminidase treatment decreased the surface-exposed sialic acids, the latter caused a serious defect in macrophage galvanotaxis without reversing the overall directionality (**Fig. 5E**), in contrast to *Salmonella* infection (**Fig. 3B**). There may be multiple glycosidases involved surface glycan modification to reverse directionality (**Fig. 5H)** as *Salmonella* possesses at least 51 putative glycosidases that likely function in glycan degradation (71). In fact, a recent study identified several glycosidases, including a putative neuraminidase, as new virulence factors essential for *Salmonella* infection of epithelial cells, which is again independent of the SPI-1 (72). We are in the process of performing genetic knockouts to identify the factors involved. It is also possible that modification of surface glycan and reduction of zeta potential were mediated by internalization during phagocytosis of the bacterium itself rather than by bacterial enzymatic activities. For instance, macrophages express Toll-like receptors (TLRs) that recognize structurally conserved molecules derived from *Salmonella* and other pathogens. All TLRs contain *N*-linked glycosylation consensus sites and both TLR2 and TLR4 require glycosylation for surface translocation and function (73, 74). Binding of *Salmonella* to these glycosylated receptors and subsequent internalization may reduce surface negativity of macrophages, leading to directional switch under EF. This idea is supported by our observations where the accumulation of certain lectin-binding aggregates within macrophages infected by *Salmonella*, but not in cells challenged by microspheres (**S8 and S9 Fig.**). It is also possible that a macrophage can uptake microspheres without significantly changing its zeta potential (*e.g*., through a neutralized receptor), and therefore still migrate to the anode.

Disseminated *Salmonella* infection is a major health problem of developing countries, responsible for ~433,000 deaths annually (75). Understanding the mechanisms that trigger dissemination is critical for efforts to target this key process for preventive and therapeutic purposes. We propose that macrophages are attracted to the site of infection by a combination of chemotaxis and galvanotaxis, driving the cells in the same direction. After phagocytosis of bacteria, surface electrical properties of the macrophage change, and galvanotaxis directs the cells away from the site of infection (**Fig. 6**). Our study represents a new perspective for the initiating mechanisms, suggesting that *Salmonella* disseminates through infection-generated bioelectrical control of macrophage trafficking. It is important to emphasize that the demonstrated bidirectional migration of macrophages to the IGEF-like electric fields is not a physical electrophoresis (movement of charged particles under direct current EF), nor through a chemical gradient that could be influenced by EF (29), but a complex, yet, poorly understood biological process that requires phosphoinositide 3-kinases and other critical signaling activities, as well as the cellular motility machinery (24, 76). Future works utilizing transgenic animals and pharmacological perturbations to target specific pathways (known or unknown) will help to pinpoint key molecules mediating the infection-dependent directional reversal (*i.e*., the molecular mechanisms to initiate disseminated infection) and response to bioelectric signaling.

**Fig. 6.**
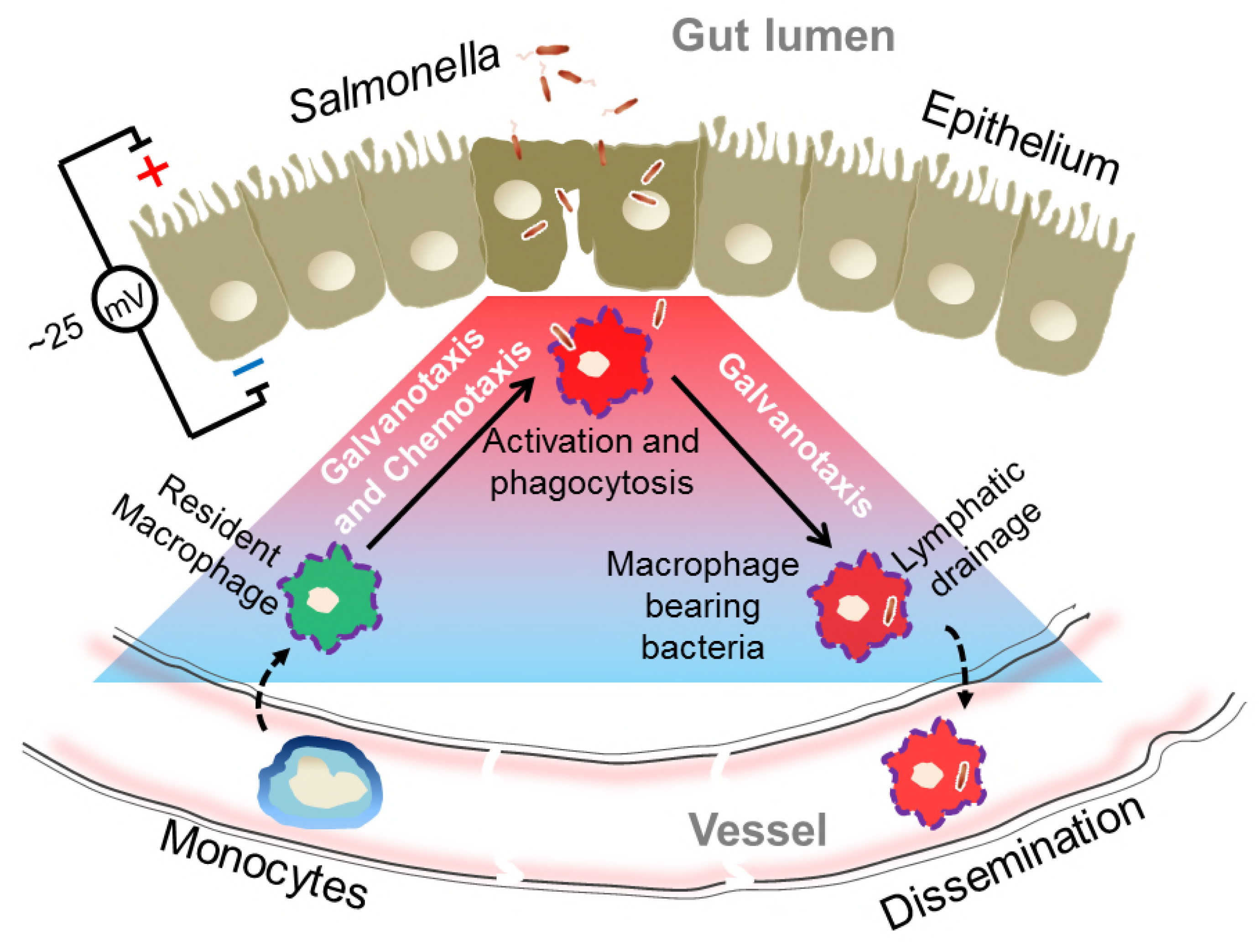
IGEF at gut epithelium and model of macrophage galvanotaxis in initiating dissemination. *Salmonella* invades and disrupts gut epithelial integrity, preferentially at the FAE, generating an IGEF (red to blue gradient) that recruits resident macrophages to the bacteria entry sites – alone or synergistically with chemotaxis. Macrophages invaded by *Salmonella* revert galvanotaxis direction through the modification of surface electric properties to reach the lymphatic drainage/blood stream, therefore initiating dissemination.

Although the present work deals primarily with gut epithelium and enteric bacteria, the general mechanism that emerged from this work could also apply to other mucous epithelia such as the respiratory tract and its associated pathogens, which is another major public health concern.

## Materials and Methods

### Mice and surgery

The mouse strains used were in a C57BL/6 background (both male and female mice were used in experiments). Mice were purchased from Jackson lab and maintained under a strict 12-h light cycle and given a regular chow diet in a specific pathogen-free facility at University of California (UC), Davis. All animal experiments were performed in accordance with regulatory guidelines and standards set by the Institutional Animal Care and Use Committee of UC Davis. In brief, we dissected mouse cecum following euthanasia, and opened longitudinally along the mesenteric attachment remnant to avoid incision damage to the single Peyer’s patch located under the antimesenteric mucosa near the apex (**S1A Fig.**). After thorough washing in mouse Ringer’s solution (154 mM NaCl, 5.6 mM KCl, 1 mM MgCl_2_, 2.2 mM CaCl_2_, 10 mM glucose and 20 mM HEPES, pH 7.4.) to remove the luminal contents, we placed the cecum with mucous side facing up, on a 30 ^o^ slope of silicone gel, prepared from polydimethylsiloxane (PDMS) in custom-made measuring chambers. The cecum was aligned and immobilized with fine metal (tungsten) pins prior to taking measurements (**S1B Fig.**). This process was usually completed within 5 min.

### Measuring transepithelial potential (TEP) with glass microelectrodes

We used glass microelectrodes to directly measure the TEP of intestinal epithelium as previously described (27, 77). TEP were recorded by microelectrode impalement through the epithelial layers. Microelectrodes (1-2 μm tip diameter; NaCl 3 M electrolyte) had resistances of ~1-2 MΩ and the potentials were offset to 0 mV prior to impalement. Cecal FAE and adjacent villus epithelium were discriminated under a dissecting microscope within a Faraday cage on an antivibration table. In some cases the TEP were measured as follows: first at the epithelial surface (0 μm), then stepwise at 50, 100 and 200 μm in depth, controlled by a micromanipulator (**S2A Fig.**). The potential typically returns to the baseline of 0 mV after microelectrode withdrawal. If the reference baseline was > ±1 and ≤ ±5 mV, the value was subtracted from the TEP recorded as shown in the equation (**S2B Fig.**); if > ±5 mV, the trace was rejected. As a control, we measured TEP of serosa epithelium and TEP of formalin-fixed mucous epithelium. Measurements were performed at room temperature in mouse Ringer’s solution. Data were acquired (saturated sampling at 100 Hz) and extracted using pClamp 10 (Molecular Devices) and analyzed using Excel (Microsoft).

### Measuring ionic currents with vibrating probes

We used non-invasive vibrating probes to measure the electric current density (J_I_) in μA cm^−2^ of mouse cecum epithelium as previously described (27, 40, 41). The probes, platinum-electroplated at the tip (~30 μm ball diameter), vibrated at a frequency between 100-200 Hz. Prior to measurements, the probe was calibrated to the experimental conditions by an applied J_I_ of 1.5 μΑ cm^−2^ (**S3C Fig.**). Under a dissecting microscope, mounted mouse ceca were positioned in the non-conductive measuring chamber (**S3A Fig.**). The plane of probe vibration was perpendicular to the epithelial surface at a distance as close as possible (**S3B Fig.**). J_I_ was recorded until the plateau peak was reached (< 1 minute) (**S3C Fig.**). Reference values were recorded with the probe away from the epithelium surface (>> 1 mm) (**S3A Fig.**). Measurements were taken at room temperature in mouse Ringer’s solution. During calibrations and measurements, a Faraday “wall” (grounded aluminum-wrapped cardboard) covered the microscope. As a control, we measured J_I_ near the surface of serosal epithelium and formalin-fixed mucous epithelium. Data were acquired and extracted using WinWCP V4 (Strathclyde Electrophysiology Software) and analyzed using Excel (Microsoft).

### Reagents, plasmids and *Salmonella* strains

Special reagents used in this work are listed in **S1 Table**. Plasmids and *Salmonella* strains used in this work are listed in **S2 Table**. Cultures of *E. coli* (for plasmid extraction) and *S*. Typhimurium were incubated aerobically at 37 °C in Luria-Bertani (LB) broth (per liter: 10 g tryptone, 5 g yeast extract, 10 g NaCl) or on LB agar plates (1.5% Difco agar) overnight. Antibiotics were used at the following concentrations unless stated otherwise: 30 μg ml^−1^ chloramphenicol, 50 μg ml^−1^ nalidixic acid, 100 μg ml^−1^ ampicillin, 50 μg ml^−1^ kanamycin and 10 μg ml^−1^ tetracycline.

### Construction of *S*. Typhimurium expressing fluorescent proteins

In an effort to monitor live intracellular *Salmonella* during macrophage galvanotaxis we constructed a *S*. Typhimurium strain derived from IR715 that constitutively expresses mCherry coded in its genome (78). In brief, the *glmS::mCherry* allele was transferred to IR715 via P22 transduction from the donor strain S. Typhimurium SL1344St (a gift from Leigh Knodler) (79), and selected by chloramphenicol. To remove the FRT-flanked chloramphenicol cassette, the transductant was transformed with pCP20 encoding FLP recombinase. The resulting strain constitutively expressing mCherry encoded by *glmS::mCherry::FRT* was designated KLL18. The GFP-expressing SPI-1 mutant Δ*invA* was generated by electroporating pGFT/RalFc into AJB75 (49). The plasmid pGFT/RalFc was constructed in two steps. First, a fragment of a *gfpmut3* gene under the control of the constitutively active kanamycin-resistance gene *aphA3* promoter was amplified from pJC43 (80) with primers of GFPmut3-F (5’-A<u>GAGCTC</u>CAGCGAACCATTTAAGGTGATAG -3’) and GFPmut3-R (5’-A<u>CTGCAG</u>TTATTTGTATAGTTCATCCATGCC -3’). This fragment was then digested with SacI/PstI and cloned into pFT/RalFc, a low-copy plasmid based on pBBR1-MCS4 (81).

### Oral infection in a mouse model of human typhoid fever

The mouse is a well-established animal model to study *Salmonella* pathogenesis (34). C57BL/6 and other mice carrying a mutation in *nramp1* develop disseminated infections when challenged by *S*. Typhimurium, which mimics human typhoid fever (82). For the mouse infection experiment, we used the hyper-disseminative D23580 strain (83, 84). In brief, D23580 was used to inoculate LB broth and incubated overnight at 37°C. C57BL/6 mice (6-10 weeks old, mixed sexes) were intragastrically infected with 10^9^ bacteria (actual inoculum was determined by plating) in 0.1 ml LB broth. Uninfected mice used as a control were given 0.1 ml sterile LB broth in place of *Salmonella*.

Mice were euthanized at the indicated time points post-inoculation by CO_2_ asphyxiation followed by cervical dislocation as the secondary method of euthanasia. Mesenteric lymph nodes (MLN) and spleens were collected aseptically and homogenized in phosphate-buffered saline (PBS) for colony-forming unit (CFU) enumeration. Ceca were dissected, cleaned, and then either mounted for bioelectrical measurement, prepared for histopathological fixation, or homogenized in PBS for CFU enumeration.

### Histology

Ceca were fixed in 10% neutral buffered formalin. After fixation, tissues were routinely processed, embedded in paraffin, sectioned and stained with hematoxylin-eosin.

### Isolation and culture of primary mouse peritoneal and bone marrow-derived macrophages

Both peritoneal and bone marrow-derived_macrophages were isolated from C57BL/6 mice (6-10 weeks old, mixed sex) following standard procedures as previously described (85). Peritoneal macrophages were seeded onto 6-well plates and allowed to adhere to the plastic for 1-2 days in DMEM (Invitrogen) with 10% Fetal Bovine Serum (Invitrogen) and 1× Antibiotic-Antimycotic solution (Invitrogen). Bone marrow-derived macrophages were cultured in the same medium as described above, but supplemented with 20% L-929 conditioned medium for 6 days (plus an extra feed at day 3), followed by one-day culture without the conditioned medium. Adherent macrophages were then harvested by gently scraping with a “policeman” cell scraper and used for subsequent experiments accordingly. Cell viability was determined by trypan blue staining and counting.

Our initial galvanotaxis experiments were carried out in peritoneal macrophages. Since we observed better directional switch in bone marrow-derived macrophages infected by *Salmonella* (**S5 Fig.**), subsequent experiments were done in bone marrow-derived macrophages, unless stated otherwise.

### Gentamycin protection assay to determine intracellular bacteria CFU

The gentamycin protection assays were carried out as previously described (86). In 24-well tissue culture-treated plates, 2×10^5^ cells were seeded per well for 5-6 h in culture medium (DMEM with 10% Fetal Bovine Serum and no antibiotics). *Salmonella* were grown overnight and used to infect macrophages at a multiplicity of infection (MOI) of 20. After 60 min of incubation, cells were gently washed 3× with PBS, and further incubated in gentamycin-containing culture media at a final concentration of 50 μg ml^−1^ for additional 60 min. Afterwards media was replaced with culture media containing 10 pg ml^−1^ gentamycin for the duration of the experiment. Intracellular CFU was measured at 16 h POI. To measure intracellular CFU, macrophages were lysed using 0.5% Tween 20 for 5 min at room temperature and released by scraping with 1 ml pipette tips. CFUs were enumerated by plating.

### Infection, challenge and treatment of macrophages

Typically, 4×10^4^ primary mouse macrophages were seeded per well of engineered silicon stencils sealed in custom-made electric field (EF) chambers (see *Engineering silicone stencil and EF chamber design* section for details) or 96-well glass bottom plates (Nunc), or 2×10^5^ cells per well in 24-well tissue culture-treated plates depending on different experiment needs, for 5-6 h in culture medium. Overnight cultures of *Salmonella* or fluorescently labeled microspheres were used to infect/challenge macrophages at a MOI of 20. The rest of the procedures were similar to the gentamycin protection assay. Cells were cultured in medium containing 10 μg ml^−1^ gentamycin for 16 h and subsequent galvanotaxis experiments were carried out in the same medium containing gentamycin.

For the neuraminidase treatment, cells were incubated in culture medium containing 100 mU ml^−1^ neuraminidase from *Vibrio cholerae* (Sigma-Aldrich) for 30 min at 37 ^o^C (50). Cells were then washed with culture medium and subsequent galvanotaxis experiments were carried out in the culture medium containing no neuraminidase.

For the low pH experiments, cells were incubated in culture medium of pH 5.8 buffered with 15 mM MES (Sigma-Aldrich) for 60 min and subsequent galvanotaxis experiments were carried out in the same media of pH 5.8 (76). Control experiments were carried out always in parallel in culture medium of pH 7.4 buffered with 14.4 mM HEPES (Invitrogen).

### Galvanotaxis assay

#### 1. Engineering silicone stencil and EF chamber design

We have tested cover glass and plastics coated with different substrates and found that macrophages perform robust and consistent galvanotaxis when cultured in tissue culture dishes (Corning). Therefore, our EF chambers were customized based on 100 mm tissue culture dishes. To facilitate group comparability and EF control, we engineered removable and reusable silicone stencils of multiple wells (diameter of 8 mm, thickness of 2.4 mm) to seed the same batch of cells that can be challenged/treated with different bacteria/substances, and monitored under identical galvanotactic conditions simultaneously (87). The EF tunnel height is fixed at around 120 μm by double-sided silicone tapes cut by a computer-controlled laser cutter (88).

#### 2. EF application and time-lapse recording

We applied exogenous EF as previously described (41, 87, 89, 90). The EF strength is based on the infection-generated EF (IGEF) we measured at the gut epithelium in two ways. First, we detected an inward J_I_ of ~1.5 μA cm^−2^ at *Salmonella-infected* FAE. The mouse Ringer’s solution we used in the measurement has a resistivity (ρ) of 19.47 mΩ cm, measured with a conductivity meter. A common approximation to the current density assumes that the current is simply proportional to the electric field, as expressed by the following equation (derived from Ohm’s law):

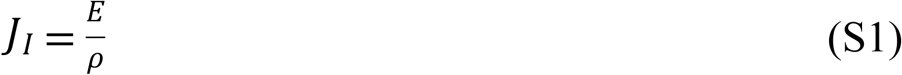

where *E* is the electric field. Plugging *J_I_* and *ρ* into the equation, we calculated that a density of 1.5 μA cm^−2^ equals an EF of 0.77 V cm^−1^. Based on trials in rabbit corneal epithelium, the *in vivo* J_I_ is likely 2-4 times larger than the *ex vivo* J_I_ because of the higher resistances of the tissues and of the higher physiological temperature. Second, we detected a TEP of ~25 mV crossing a single layer of *Salmonella-infected* gut epithelium. This generates an EF of 5 V cm^−1^ providing an epithelial cell height of 50 μm. After testing a range around those values, we have empirically chosen an EF of 4 V cm^−1^ as it consistently induced significant directional migration of primary mouse macrophages, although biased directional migration can be achieved by an EF as small as 0.5 V cm^−1^. Actual EF strengths were measured and determined with a voltmeter before and after each EF application.

Cell migration was monitored with a Carl Zeiss Observer Z1 inverted microscope equipped with a motorized stage and an incubation chamber (37 ^o^C and 5% CO_2_). Time-lapse contrast images and/or images of appropriate fluorescence channels were captured using MetaMorph NX program (Molecular Devices). A Retiga R6 (QImaging) scientific CCD camera and long exposure time (~2 s) were used to detect and monitor intracellular *Salmonella* expressing fluorescent proteins. Typically, in each experiment, 2-4 fields of each condition under a 10× or a long distance 20× lens were chosen. Images were taken at 5 min intervals for up to 3 h unless stated otherwise.

#### 3. Image process and data analysis/presentation

Time-lapse images were imported, processed and assembled in ImageJ (http://rsbweb.nih.gov/ij). To quantify single-cell and population motility we extracted the trajectory of each cell migration (> 30 cells for each condition) using an automatic/manual tracking tool (41, 76, 87). Directionality as directedness in cosine theta (*θ*), where *θ* is the angle that each cell moved with respect to the EF vector, were quantified from the coordinates of each trajectory (91, 92). If a cell moved perfectly along the field vector toward the cathode, the cosine of this angle would be 1; if the cell moved perpendicular to the field vector, the cosine of this angle would be 0; and if the cell moved directly toward the anode, the cosine of this angle would be -1. Dead cells irresponsive to EFs were either washed away or excluded from quantification by thresholding. The galvanotaxis assays and quantification of directionality in bone marrow-derived macrophages infected with Δ*invA* and their isogenic wildtype *Salmonella* were assigned in a double-blinded manner.

To simulate cell migration, each cell was numbered and its *x* and *y* coordinates were measured on the first image and on every subsequent image in the image stack, with the x-axis parallel to the applied electric field. The (*x, y*) data of each cell was imported with the ImageJ chemotaxis tool plugin, and recalculated based on the optical parameters (lens and camera). Trajectories of the cells in each group were simulated in a Cartesian coordinate system by placing the first coordinates of each cell in the origin (0, 0).

To plot the rose histograms, we combined *θ* of each time interval of tracked cells in each group. The vector *θ*, expressed in radians, were calculated from the coordinates of each trajectory. The distribution of *θ* in 12 angle bins and their abundance in percentage were plotted in Matlab (Mathworks) using a custom script (available upon request).

#### 4. Morphological analysis

The polarity of macrophages was determined by the relative distribution of the characterized protrusive lamellipodia front and uropod tail with respect to the applied EF. These were done by visually inspecting a large number of cells (> 50 cells in each case) from images taken at 3 h after EF exposure or by quantifying cellular actin intensity of confocal images using ImageJ software with line scan and color function plugins (87).

### Immunobiochemistry, lectin staining and confocal microscopy

Macrophages were seeded in either 96-well glass bottom plates (Nunc) or in custom-made EF chambers, and infected/challenged/treated by following procedures as described above. The cells were fixed with 4% paraformaldehyde immediately or after EF exposure for 3 h with field orientation marked. *Salmonella* were detected with a polyclonal antibody specific to *Salmonella spp*. (Mybiosource) stained by an Alexa Fluor 488-conjugated secondary antibody. F-actin was labeled by Alexa Fluor 555 Phalloidin. Nuclei were labeled by Hoechst 33342.

In the cases of lectin staining, fixed cells were incubated with FITC-labeled lectin (**S3 Table**) overnight at 4 ^o^C, washed extensively and then stained with DAPI (life technology) for 10 min on ice.

Cells were photographed using either an inverted (for cells on cover glass with no EF) or an upright (for cells on plastic EF chambers) Leica TCS SP8 confocal microscope (Leica microsystem). Images were processed using ImageJ. Quantification and comparison of fluorescent intensity were done in images taken in the same batch with the same optical setup and parameters. Lectin binding aggregates stained after permeabilization, were quantified by thresholding. Cells were counted using particle analysis function.

### Flow cytometry

Infected/challenged/treated macrophages, handled according to the procedures described above, were then incubated with Fc-block (BD) on ice for 15 min and stained with FITC-labeled lectin (**S3 Table**) for 1 h on ice and then stained with Aqua-live/dead (life technology) for 30 min at room temperature. Cells were washed after each step and before being analyzed on a BD Fortessa flow cytometer. Data were analyzed using FlowJo software (Tree Star Inc.). After gating single cells and live cells, the geometric mean fluorescence intensity and standard error were collected for each FITC lectin in each condition in addition to FMO (Fluorescence Minus One) for FITC-Lectin (no FITC-lectin staining). The geometric mean fluorescence intensity was then standardized across experiments using the following equation:

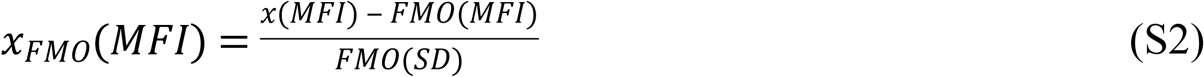

where *x_FMO_*(*MFI*) is the standardized geometric mean fluorescence intensity of a specific lectin for a specific experiment, *x*(*MFI*) is the geometric mean fluorescence intensity of a specific lectin for a specific experiment, *FMO*(*MFI*) is the geometric mean fluorescence intensity of the FMO for a specific experiment and *FMO*(*SD*) is the standard deviation of the FMO for a specific experiment. Standardized geometric mean fluorescent intensities were then plotted and tested for statistical significance (**S10 Fig.**).

### Measuring zeta potential

Macrophages were seeded onto 24-well tissue culture plates (Nunc) and infected/challenged/treated following procedures as described above. Cells were fixed in 2% paraformaldehyde and washed with motility buffer (10^−4^ M potassium phosphate buffer at pH 7.0, with 10^−4^ M EDTA) (64). Cells were then gently collected by scraping with a “policeman” cell scraper and subsequent measurements were done in motility buffer, except for the macrophages tested for acidic treatment, which were measured in either pH 5.8 medium or pH 7.4 medium as a control. Zeta potential was determined by electrophoretic light scattering at 25 °C with a Zetasizer (Malvern). Zeta potential was calculated in mV and differences between groups were analyzed by Student’s *t*-test.

### Statistics

Galvanotaxis data from representatives of at least four independent experiments were routinely presented as mean ± standard error, unless stated otherwise. Student’s *t*-test was used for paired or unpaired comparisons. Distributions of macrophage polarity between control and neuraminidase treated or between neutral and acidic conditions were analyzed using χ^2^ test. One-way ANOVA and Student’s *t*-test were used for comparisons of standardized geometric mean lectin fluorescent intensity among one or two or more groups, respectively, using Excel (Microsoft). ns: non-significant, * *P* < 0.05, ** *P* < 0.01, *** *P* < 0.001.

## Acknowledgments

We are deeply grateful to R. M. Tsolis and A. J. Bäumler for providing the *Salmonella* strains and hosting part of the animal works. We thank S. Hwang and C. Bevins for critical suggestions; V. E. Diaz-Ochoa, B. Young, L. Olney, D. Duo and I. Brust-Mascher for technical assistance; L. Knodler for sharing a *S*. Typhimurium SL1344 constitutively expressing mCherry; and Robert Heyderman for providing *S*. Typhimurium strain D23580. This work was supported by US Army Research Office grant W911NF-17-1-0417 to A. M.; and an inter-department seed grant S-MPIDRGR from UC Davis, SOM to M. Z., R. M. Tsolis. and Y. H. S. F. F. was supported by Fundação para a Ciência e a Tecnologia (SFRH/BD/87256/2012). E. M. M. is supported by an early career award from the Burroughs Wellcome Fund and by NIH 1DP2OD008752.

## Supplementary Information for

This PDF file includes:

S1 to S11 Figures

S1 to S3 Tables

Captions for S1 to S5 Movies

Other Supplementary Materials for this manuscript includes the following:

S1 to S5 Movies

**S1 Fig.**
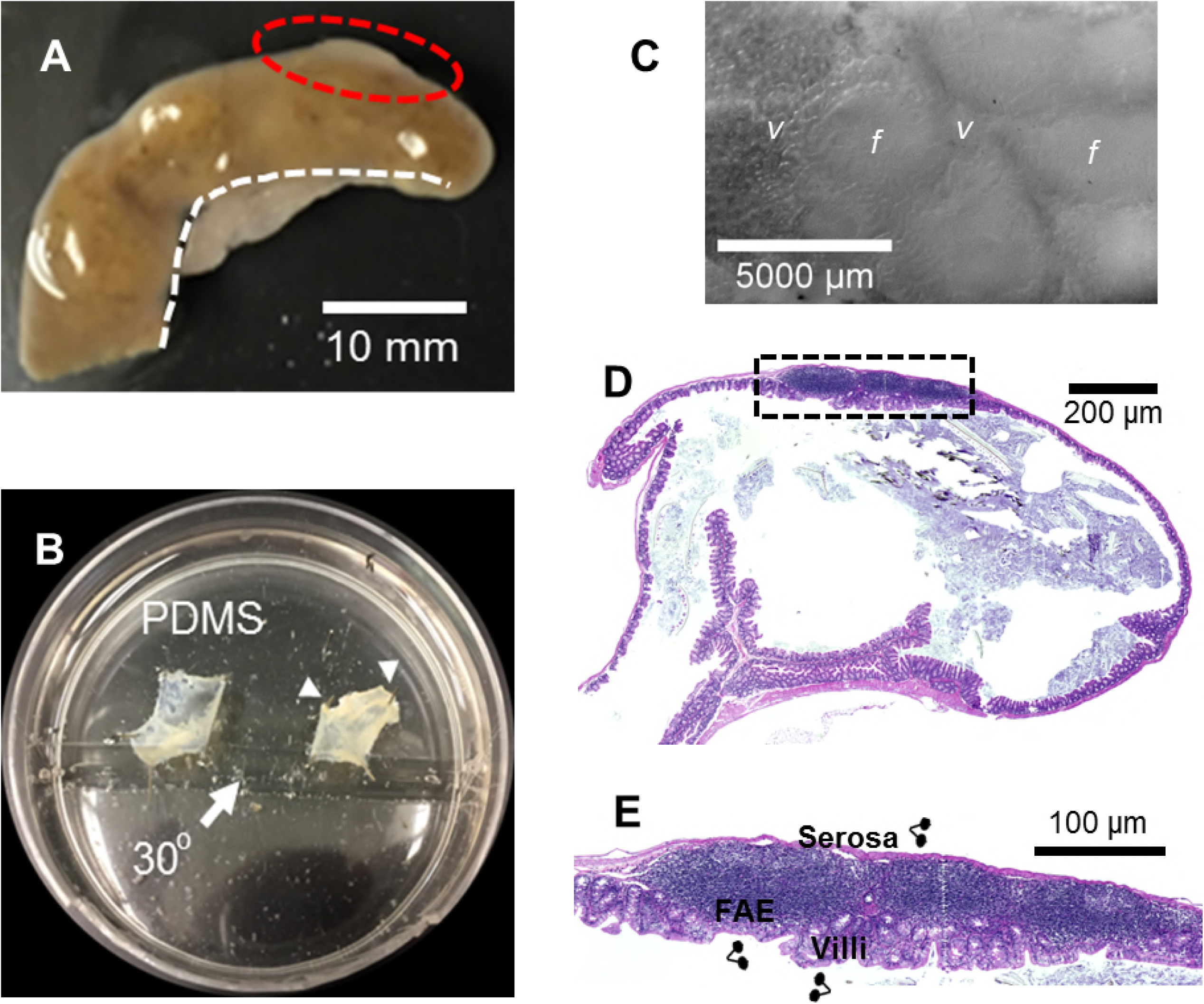
An *ex vivo* mouse cecum model for characterization of bioelectrical activities of gut epithelium. (A) Cecum was dissected from C57BL/6 mouse and opened longitudinally along the mesenteric attachment remnant (dotted white line) to avoid incision damage of lymphatic structure that is located under the antimesenteric mucosa near the apex (red circle). (B) Cecum mounted in a custom-made chamber with mucous epithelium facing up on a 30 ^o^ slope of silicone gel and held with the cecum edge with fine metal pins (white arrow heads). (C) Mouse cecal epithelium under a dissecting microscope. FAE -- the smooth appearing regions (f), and inter-follicle/surrounding villi (v) are shown. Bar, 5000 μm. (D) Hematoxylin and Eosin (H&E) stain of a mouse cecum showing the structure of a Peyer’s patch. Bar, 200 μm. (E) Magnification of the checked area of panel D showing FAE and inter-follicle and surrounding villi. Double-dotted forks indicate the sites where TEP and J_I_ were measured. Serosal epithelia were served as controls. Bar, 100 μm.

**S2 Fig.**
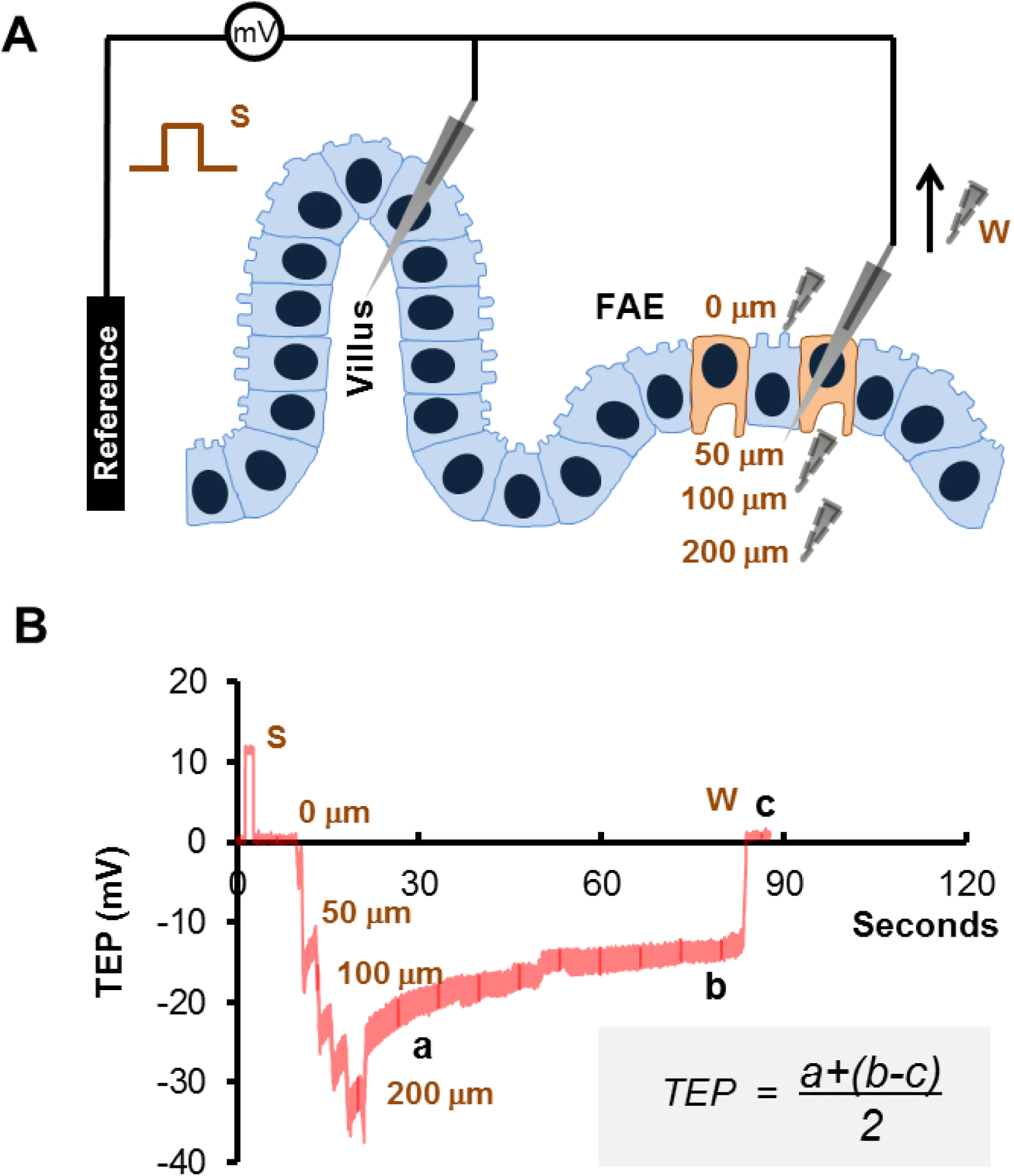
Measuring TEP with microelectrodes. (A) Schematic depicting the microelectrode setup, sites and procedures of measurement. The measuring microelectrode was impaled through the FAE or surrounding villus epithelial layer (one site at a time) and the circuit was closed by a reference electrode placed in the buffer, representing the lumen. Microelectrode resistance (S, where 10 mV equates 1 MΩ) was generated and recorded prior to each impalement to ensure that the tip was neither broken nor obstructed. In some cases the TEP of infected FAE were measured as follows: first at the epithelial surface (0 μm), then in stepwise at 50, 100 and 200 μm in depth. The potential typically returns to the baseline of 0 mV after microelectrode withdrawal (W). (B) A representative result as shown in Fig. 1C, and a specific equation used to calculate TEP value as a modified mean from raw data. In the equation, ‘a’ and ‘b’ are the early and late values of each impalement where the electrode was kept in position for at least 60 seconds. ‘c’ is the reference value immediately after the electrode withdrawal.

**S3 Fig.**
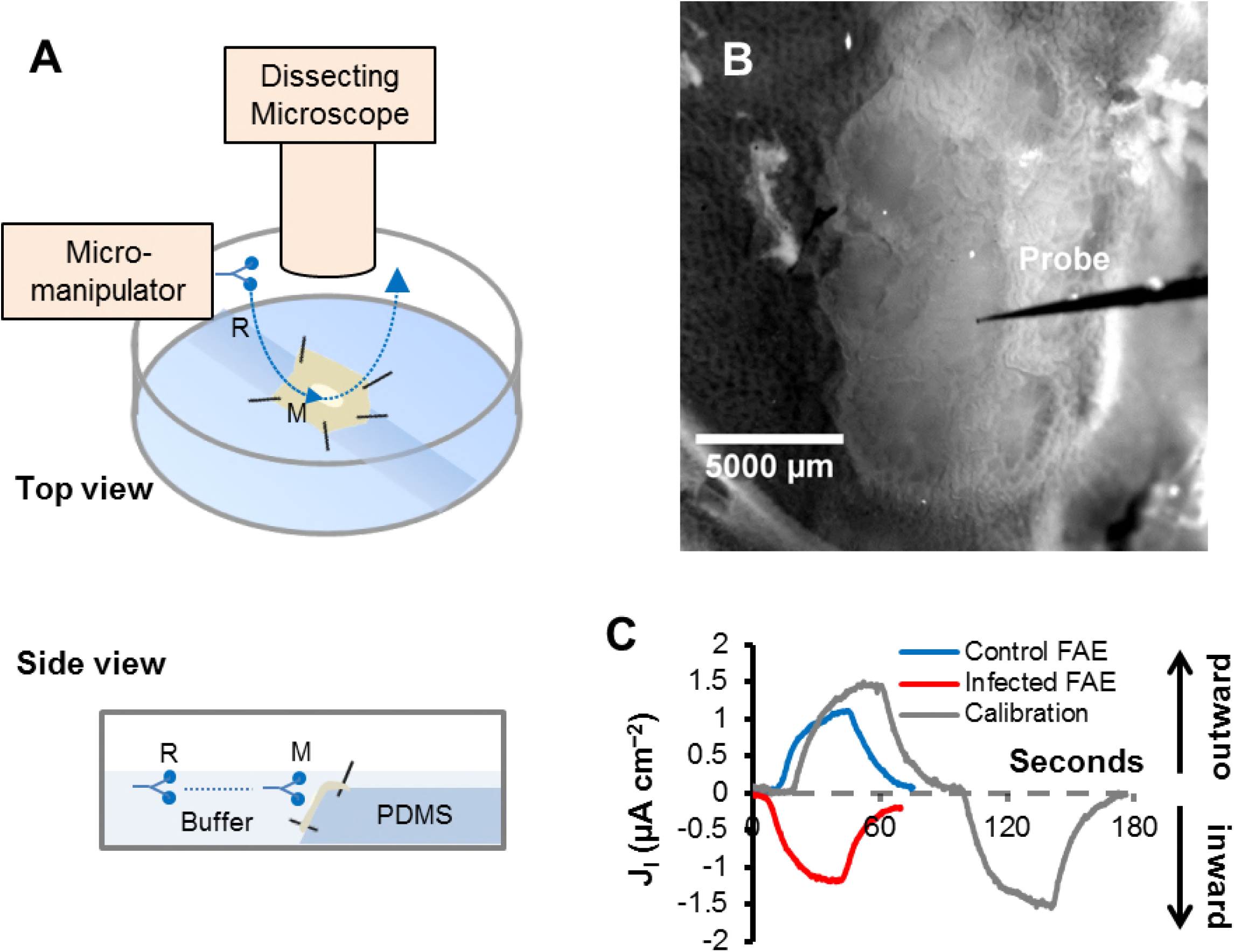
Measuring bioelectric currents using vibrating probes. **(A)** Schematic of equipment setup and measuring procedures. Under a dissecting microscope, a probe vibrating between 100-200 Hz controlled by a micromanipulator is moved from the reference position (R) to the measuring position (M), as close as possible (~20 μm) to the epithelial surface, to detect ionic currents. **(B)** A picture of a vibrating probe approaching an FAE. **(C)** Representative results of bioelectric current measured at FAE from Salmonella-infected or control mouse cecum. Probes were calibrated by passing a 1.5 μA cm^−2^ electric current through the measuring buffer in either direction. By convention, flux of positive charge is used for electric current direction. As in most studies, we used conventional current flow, thus outward current density is defined as net positive charge leaving the epithelial surface and inward current densities as that entering. Hence, positive values represent net outward current densities and negative values represent net inward current densities.

**S4 Fig.**
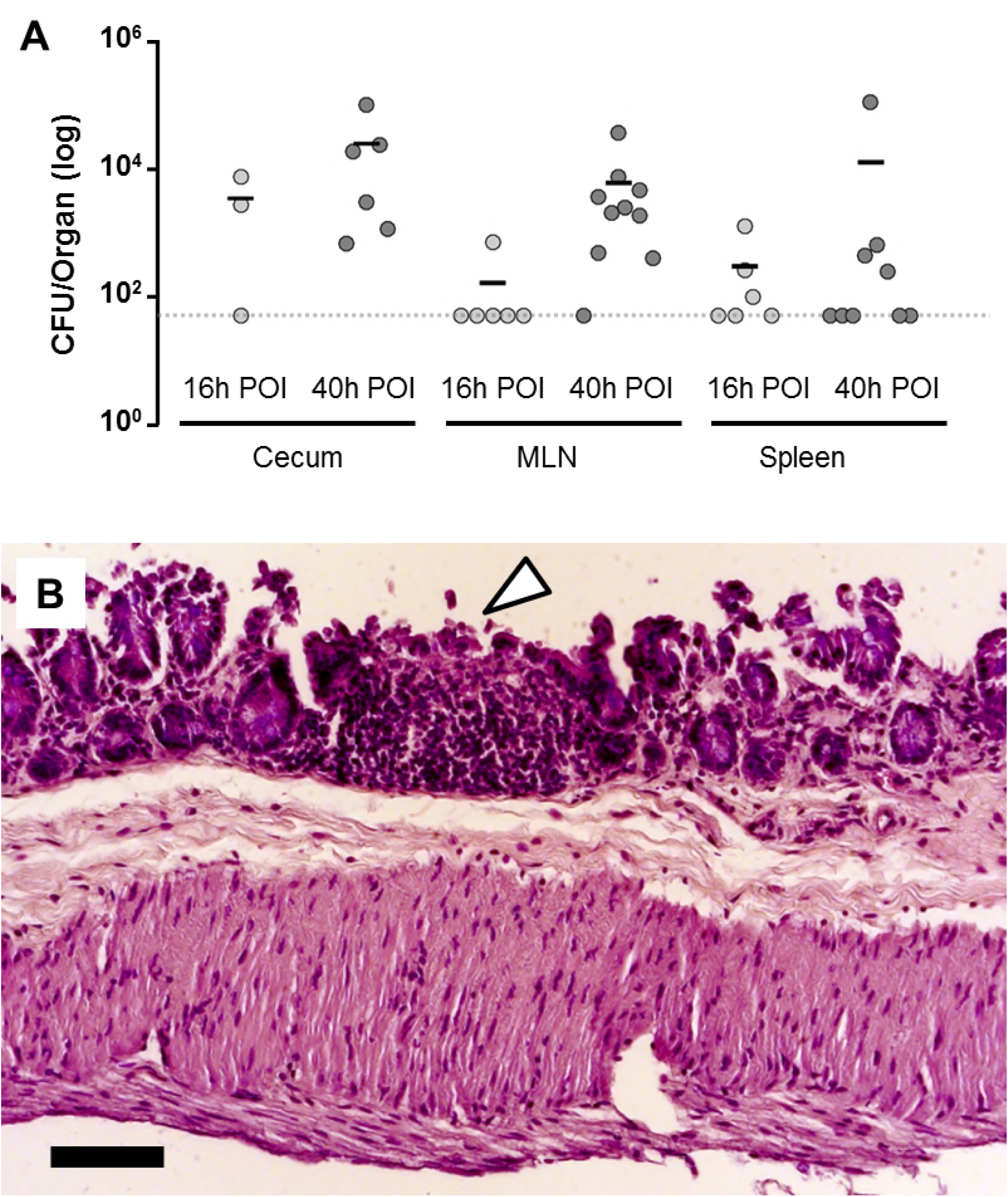
Invasion of cecal epithelium and early dissemination. (A) Ceca, free of contents, mesenteric lymph nodes (MLN) and spleens were dissected in sterile conditions at 16 h or 40 h post of infection (POI) from mice orally infected with *S*. Typhimurium. Bacterial loads were determined by homogenizing each specimen, serial dilution and plating on LB plates. Colony forming units (CFU) were calculated by counting bacterial colonies selected with appropriate antibiotics. CFUs lower than the detection limit as indicated by the dotted line, were treated as the limit. (B) H&E stain of a cecum section from mice orally infected with *S*. Typhimurium. Disruption of the follicle associated epithelium (arrowhead) and thickened intestinal wall are shown. Bar, 100 μm.

**S5 Fig.**
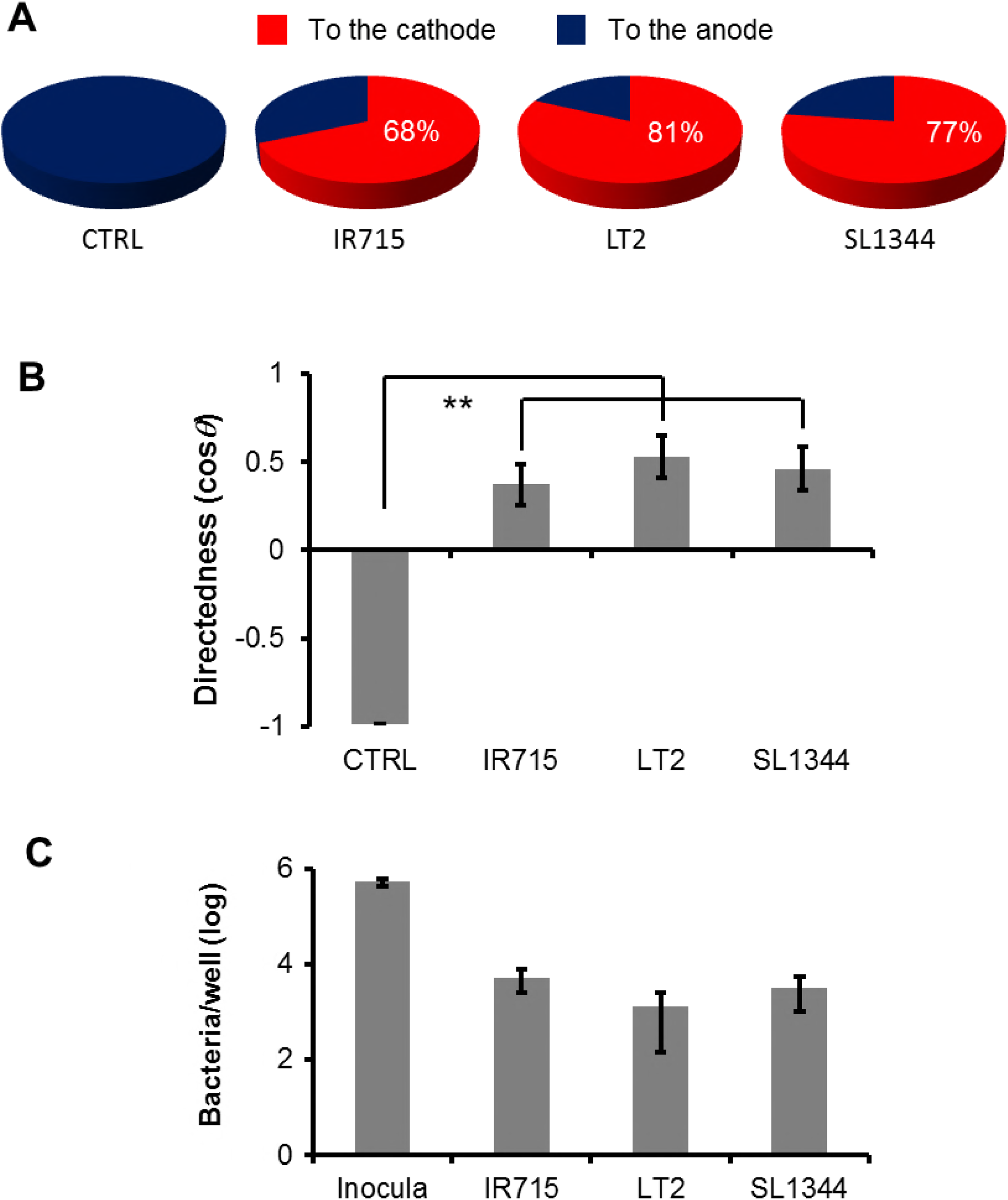
Robust cathodal direction switch in bone marrow-derived macrophages infected with different virulent *Salmonella* strains. Bone marrow-derived mouse macrophages challenged with three virulent *S*. Typhimurium at a MOI of 20, subject to galvanotaxis assay at 16 h POI. (A) Pie charts show percentage of cells migrating to the cathode (red) or to the anode (blue). (B) Overall directionality. ** *P* < 0.01 by unpaired Student’s *t*-test. (C) Quantification of intracellular bacteria. Bone marrow-derived macrophages were seeded in 24-well plates and challenged with *S*. Typhimurium at an MOI of 20. Actual inocula were determined by plating and colony counting. Intracellular bacteria at 16 h POI was determined by a gentamycin protection assay. Representative data present as log CFU per well, normalized to each inoculum. Bar in S.E. from triplicate wells.

**S6 Fig.**
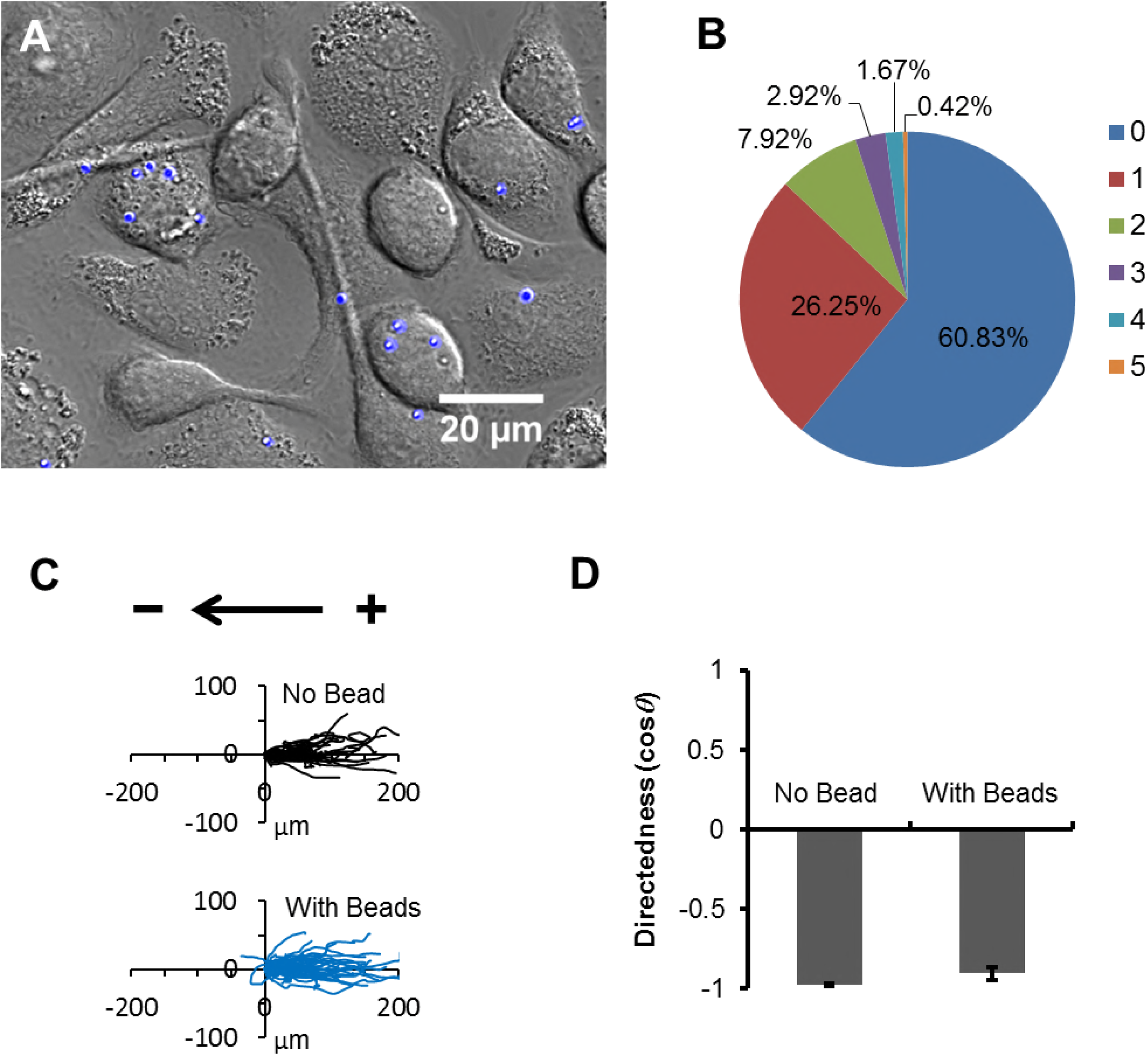
Phagocytosis (of microspheres) *per se* does not affect directional migration of macrophages in response to EF. (A) Macrophages were seeded in 96-well glass-bottom plates and challenged with 1 μm microspheres with blue fluorescent at a MOI of 20. Excessive microspheres were removed by washing with medium. At 16 h POI cells bearing beads (blue) were counted under an epifluorescence microscope. (B) Pie chart of percentage of cells with zero, one or up to 5 bead(s) in a typical experiment. (C) Trajectories and (D) directedness of macrophages under an EF of 4 V cm^−1^ in the indicated orientation for 3 h.

**S7 Fig.**
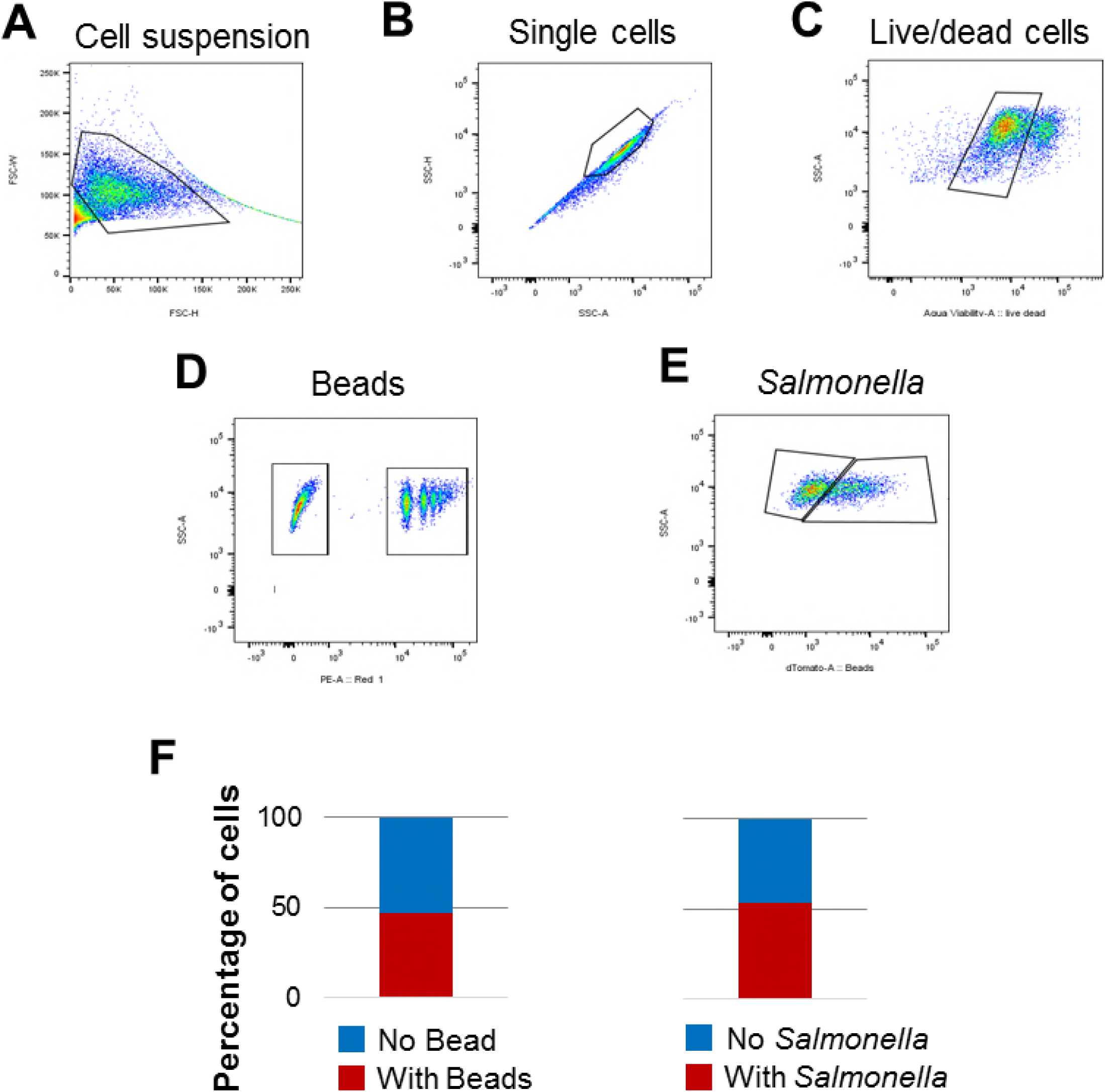
Determining challenge/infection rates by flow cytometry. Macrophages were challenged with 1 μm, red fluorescently labeled polystyrene microspheres or live *S*. Typhimurium IR715 constitutively expressing mCherry at an MOI of 20. Excessive beads and bacteria were removed by washing. Residual extracellular bacteria were killed per gentamycin treatment. (A-E) Representative flow cytograms of a complete experimental design to count target cells using flow cytometry. (A) At 16 h POI cells were harvested and labeled with Aqua blue and analyzed by flow cytometry. (B) An example of gates used to exclude fragments and cell clumps. (C) Dead cells were excluded by gating Aqua blue signal. Live cells were subject to further cell counting in either PE fluorescence channel for red fluorescent beads (D) or with the dTomato fluorescence channel for *Salmonella* expressing mCherry (E). (F) Representative bar charts showing percentage of macrophages containing intracellular bacteria or beads.

**S8 Fig.**
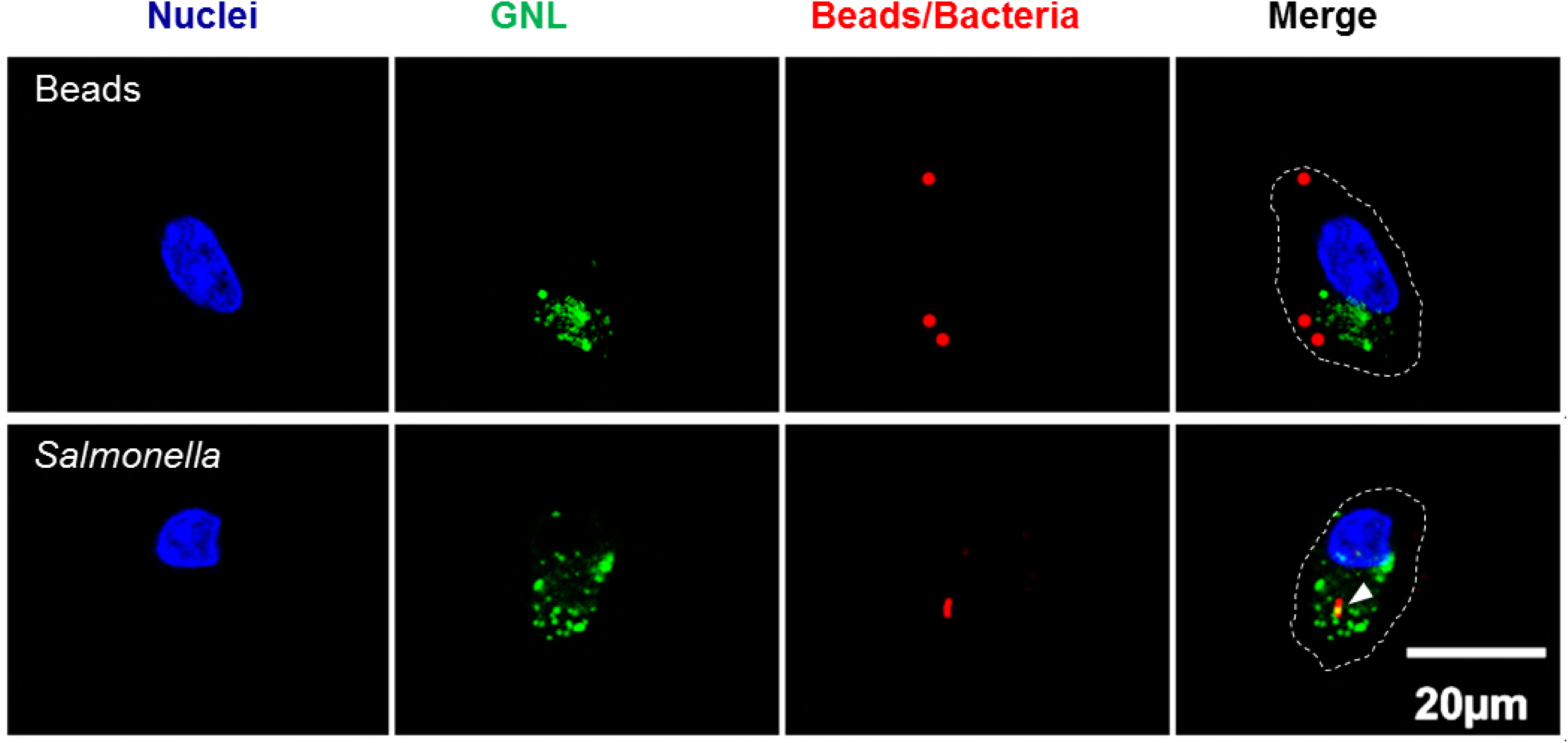
GNL-binding aggregates in macrophages infected with *Salmonella*. Representative confocal photographs show macrophages containing red fluorescence-labeled beads or *Salmonella* expressing mCherry (red). Cells were fixed, permeabilized and stained with DAPI (blue) and FITC-conjugated GNL (green). Cells were outlined in merged photographs (white dashed line). Bar, 20 μm. Note the GNL-binding aggregates inside macrophages containing intracellular *Salmonella* (white arrowhead).

**S9 Fig.**
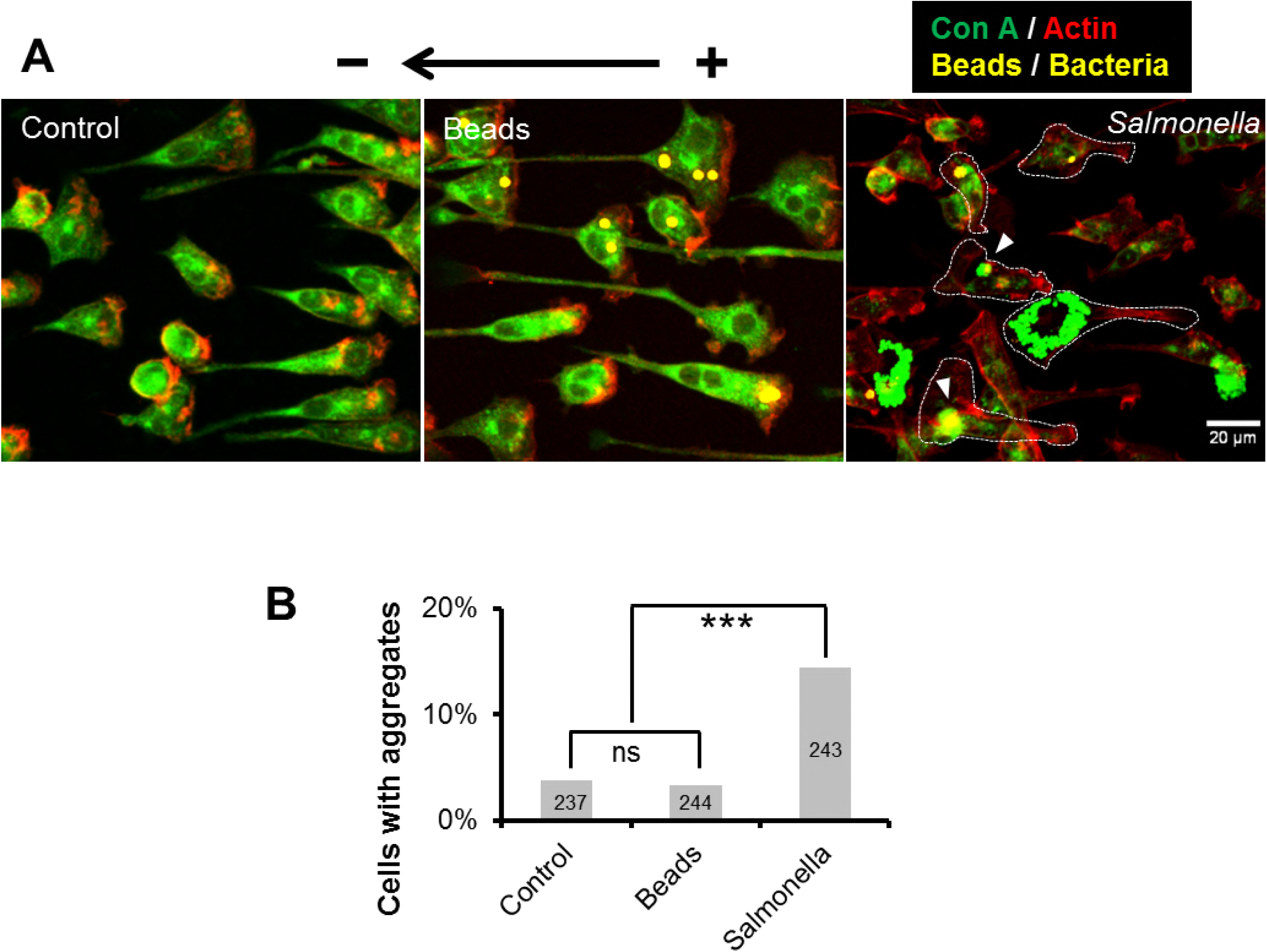
Con A-binding aggregates in macrophages infected by *Salmonella*. (A) Galvanotaxis assays were performed per the rigorous experiment design illustrated in Fig. 3A. Cells were fixed, permeabilized and labeled with Alexa Fluor 555 Phalloidin (red) and FITC-conjugated Con A (green), and scanned with an upright confocal microscope. Phagocytosed beads and intracellular bacteria were pseudocolored in yellow. Bar, 20 μm. Control macrophages (left panel) and macrophages challenged with beads (middle panel) were exclusively polarized to the anode with a characteristic morphology: massive actin meshwork in the front and a uropod at the rear. Cells infected with *Salmonella* (right panels) reversed their polarity to the cathode. Note the significant Con A-binding aggregates in macrophages containing intracellular *Salmonella* (white arrowheads). (B) Quantification of macrophages with Con A aggregates. Counted cell numbers are indicated inside each bar. *** *P* < 0.001 by *χ*^2^ test.

**S10 Fig.**
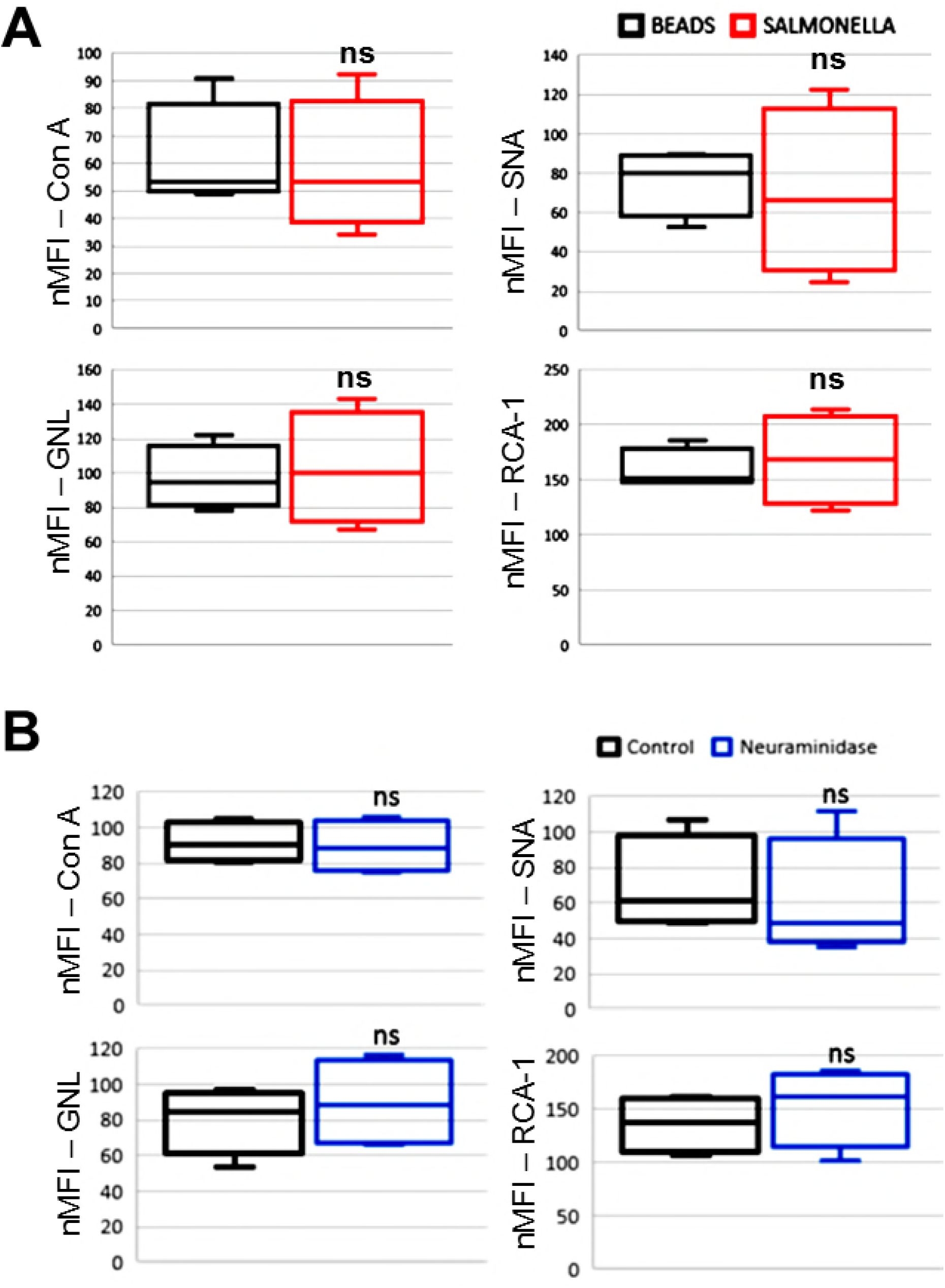
The effects of *Salmonella* infection and neuraminidase treatment on the binding of selected lectins. Box plots showing normalized mean fluorescence intensity (nMFI) of macrophages, either (A) challenged with microspheres and *Salmonella* at 16 h POI, or (B) incubated with 0 or 100 mU ml^−1^ neuraminidase for 30 min. Cells were stained with Con A, SNA, GNL or RCA-1, and analyzed by flow cytometry. Data from 4 independent experiments. ns: non-significant by unpaired Student’s *t*-test.

**S11 Fig.**
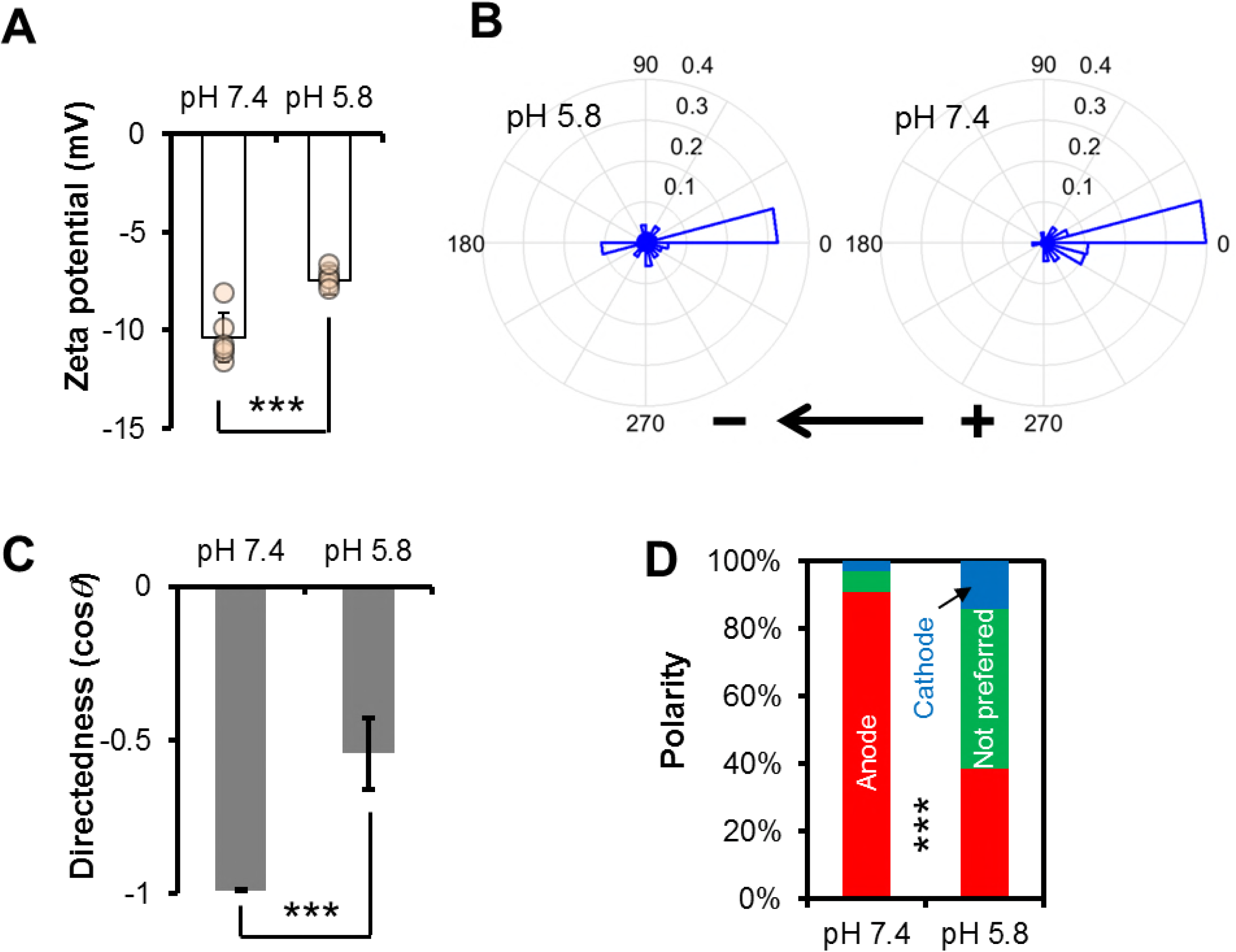
Low pH reduces surface charge and impairs macrophage galvanotaxis. (A) Zeta potential of macrophages cultured in pH 7.4 or pH 5.8. *** *P* < 0.001 by Student’s *t*-test. (B) Rose plots and (C) directedness of macrophages cultured in pH 7.4 or pH 5.8 exposed to an EF of 4 V cm^−1^. *** *P* < 0.001 by Student’s *t*-test. (D) Polarity of macrophages cultured in pH 7.4 or pH 5.8, exposed to an EF of 4 V cm^−1^ for 3 h. Data was quantified from a representative of two independent experiments. *n* = 65 and 78 for macrophages cultured in pH 7.4 or pH 5.8, respectively. *** *P* < 0.001 by *χ*^2^ test. See also Movie S5.

**S1 Table:** Special reagents used in this study.

| Reagent | Description | Source |
| --- | --- | --- |
| Microspheres | FluoSpheres® Carboxylate-Modified Microspheres, 1.0 µm, blue fluorescent (365/415), 2% solids | Invitrogen, F8814 |
| Microspheres | FluoSpheres™ Carboxylate-Modified Microspheres, 1.0 µm, red fluorescent (580/605), 2% solids | Invitrogen, F8821 |
| Neuraminidase | Neuraminidase (Sialidase) from *Vibrio cholerae* | Roche, 11080725001 |
| Anti-*Salmonella* | Polyclonal antibody | Mybiosource, MBS535017 |
| Phalloidin | Alexa Fluor™ 555 Phalloidin | Invitrogen, A34055 |
| Streptavidin | Fluorescein Streptavidin | Vectorlabs, SA-5001 |
| Aqua blue | LIVE/DEAD™ Fixable Aqua Dead Cell Stain Kit, for 405 nm excitation | Invitrogen, L34957 |

**S2 Table:**
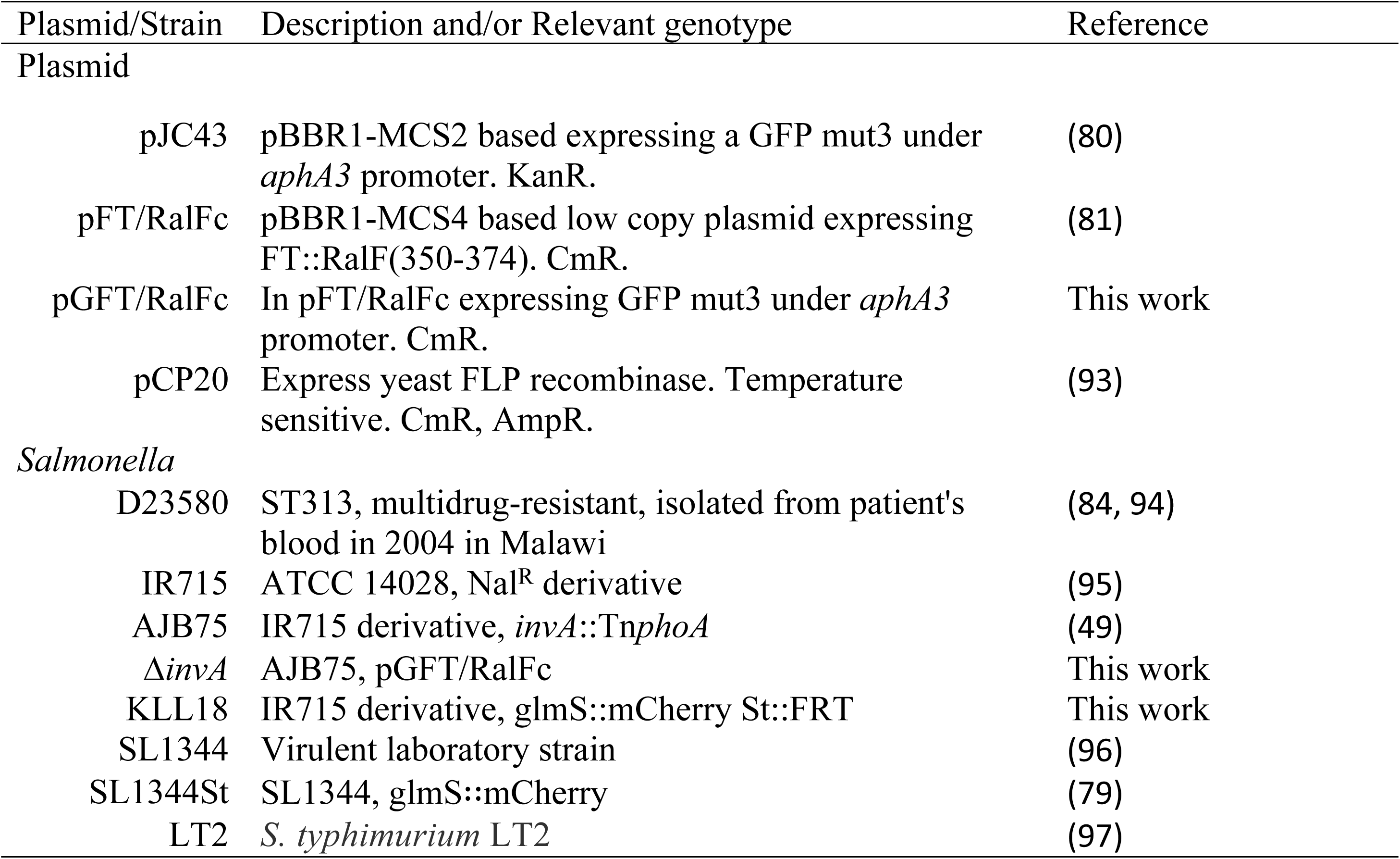
Plasmids and *Salmonella* strains used in this study.

**S3 Table:** Selected lectins used in this study.

| Lectin | Description | Sugar specificity | Source |
| --- | --- | --- | --- |
| GNL | Fluorescein labeled<br>Galanthus Nivalis Lectin | α-Mannose | Vectorlabs, FL-1241 |
| Con A | Fluorescein Labeled<br>Concanavalin A | α-Mannose, D-Glucose | Vectorlabs, FL-1001 |
| SNA | Fluorescein labeled<br>Sambucus Nigra Lectin | N-Acetylgalactosamine | Vectorlabs, FL-1301 |
| RCA-1 | Fluorescein labeled Ricinus<br>Communis Agglutinin I | D-Galactose | Vectorlabs, FL-1081 |
| MAL-2 | Biotinylated Maackia<br>Amurensis Lectin II | N-Acetylneuraminic acid<br>(sialic acid) | Vectorlabs, B-1265 |

### S1 Movie

Primary mouse peritoneal macrophages migrate to the anode in response to an EF of 4 V cm^−1^ in the indicated orientation for 5 h 50 min. Time-lapse phase contrast images were acquired one frame every 5 min. Bar, 50 μm.

### S2 Movie

Unidirectional migration of macrophages to the anode in control (top) (trajectories in highlighted yellow line) and bidirectional migration of macrophages challenged with *S*. Typhimurium (bottom), either to the cathode (trajectories in highlighted red line) or to the anode (trajectories in highlighted yellow line), under an EF of 4 V cm^−1^ in the indicated orientation for 3 h. Time-lapse phase contrast images were acquired one frame every 5 min. Bar, 50 μm.

### S3 Movie

Galvanotaxis of control macrophages (top left) or macrophages challenged with microspheres (bottom left) or wildtype *Salmonella* (bottom right) or SPI-1 mutant Δ*invA* (top right), under identical conditions at an EF of 4 V cm^−1^ in the indicated orientation for 3 h. Time-lapse phase contrast and fluorescence images were acquired one frame every 5 min sequentially, and combined using ImageJ. Note the opposite directional migration of macrophages containing beads (blue) or *Salmonella* (red and green), to the anode or to the cathode respectively. Bar, 50 μm.

### S4 Movie

Impaired directional migration of macrophages treated with neuraminidase (left), compared in parallel to the unidirectional migration of control macrophages (right) to the anode under an EF of 4 V cm^−1^ in the indicated orientation for 2 h 45 min. Time-lapse phase contrast images were acquired one frame every 5 min sequentially and combined by using ImageJ. Bar, 100 μm.

### S5 Movie

Galvanotactic behaviors of macrophages cultured in acidic condition of pH 5.8 (top) compared, in parallel, to the unidirectional migration of macrophages cultured in pH 7.4 (bottom) to the anode, under an EF of 4 V cm^−1^ in the indicated orientation for 3 h. Time-lapse phase contrast images were acquired one frame every 5 min sequentially and combined using ImageJ. Bar, 100 μm.

